# Deconstructing “Boldness”: Latent behavioural structure across novelty, predation, and exploration in an endangered species translocation program

**DOI:** 10.64898/2026.11.10.765630

**Authors:** Laney H. Nute, Richard Buchholz

## Abstract

Risk-related behaviours including neophobia, exploration, and predator avoidance are often assumed to covary along a general “boldness” axis. Although some translocation studies have linked one or more of these behaviours with fitness, it is possible that the underlying behavioural structure varies under different selective environments. Conservation breeding programs provide a unique opportunity to evaluate the structure of behavioural correlations in a non-domestic species under artificial conditions. We tested whether risk-related behaviours represent a unified boldness syndrome in a captive population of the endangered Attwater’s prairie chicken (APC; *Tympanuchus cupido attwateri*). We quantified the repeatable behaviours of 257 juveniles and compared 11 models of behavioural covariation structure using multivariate mixed models and confirmatory factor analysis. There was no support for a single risk-taking axis; instead, we found three dimensions representing responses to novelty and environmental stimuli (Engagement), predator avoidance (Threat Response), and maze exploration (Maze Progression). Our results indicate that these risk-related behaviours are not organized along a single axis in captive APCs, challenging assumptions that risk-taking represents a universal behavioural syndrome. To anticipate post-release success accurately, conservation managers should understand the exact behavioural architecture of the captive population rather than assume that the classical bold-shy structure occurs in all contexts.

## Introduction

Behavioural types are repeatable behavioural differences among individuals, while correlations among these behaviours constitute behavioural syndrome structures that may constrain adaptive responses to selection (1). Considerable work examining the heritability and fitness consequences of behavioural syndromes have demonstrated that behavioural consistency is influenced by both genetic architecture and environmental conditions (2–4). Additive genetic correlations underlying behavioural syndromes result in a 33% constriction of a population’s ability to respond to selection (5). Selection acting on one behavioural trait can indirectly alter others when they are genetically correlated, as observed for the exploratory and risk-taking behaviours during selection experiments in great tits (*Parus major)* (6). The direction of correlation between traits, however, is not always stable. In laboratory mice (*Mus musculus*) strains selectively bred for increased wheel-running behaviour, offspring from crossed lines exhibited increased running speed but decreased running duration, indicating that similar behavioural phenotypic outcomes can emerge through the evolution of different underlying trait combinations (7). Behavioural correlations can also degrade if environments change. Research has shown that the correlation between risk-taking and aggression strengthen under high predation pressure but weaken or disappear in low-risk environments (8–10). Together, these findings suggest that behavioural syndromes are dynamic structures whose integration may shift depending on ecological or selective conditions.

The dynamic structures of behavioural syndromes have motivated calls to determine whether the traits commonly grouped within a syndrome truly represent a unified functional axis (11). Among the most widely studied behavioural syndromes is boldness, broadly defined as the propensity to take risks (12–15). As risk-taking behaviour can mediate trade-offs between resource acquisition and mortality risk, it is frequently assumed to represent a key axis of variation on which natural selection can act. Measures of the boldness axis have included neophobic reactions, threat responses such as predator avoidance behaviour, and exploration tendencies, and have been demonstrated to affect survival (16–20). Although these behaviours are sometimes treated as interchangeable indicators of a generalized risk-taking phenotype, selection may act differently on each component depending on the environmental context (13,21).

Captive breeding programs provide a particularly relevant system for evaluating how altered selective environments shape the structure of behavioural syndromes. In captivity, a lack of predators may relax selection on antipredator responses (22), while novel conditions including human exposure, artificial rearing environments, and management practices may impose new forms of selection on exploratory and risk-related behaviours (23–25). Additionally, captive populations are often subject to intensive genetic management designed to preserve genetic diversity, potentially altering not only trait means but also covariance among behavioural traits. These processes raise the possibility that behaviours commonly interpreted as components of a unified “risk-taking” syndrome may become decoupled under captive conditions.

To test whether the predicted integration among risk-taking measures is evident under altered selective pressures in captivity, we evaluated the behavioural structure of risk-taking in a captive breeding population of the critically endangered Attwater’s prairie chicken (APC; *Tympanuchus cupido attwateri*). Specifically, we tested a set of *a priori* models (Table 1; electronic supplementary material, Fig. S1-S3) that predicted whether common measures of boldness including neophobia, antipredator behaviour, and exploration would represent semi-independent traits or correlated components of a larger risk-taking syndrome. These behavioural contexts are particularly relevant to the APC beyond the abrupt change in environments that occurs during release. Due to anthropogenic alteration of critical habitats (26), APCs are forced to explore novel environments as they arise while facing predation from both terrestrial and aerial predators. Without comparable data from wild conspecifics, our study cannot test whether captivity has altered behavioural relationships relative to wild conspecifics. Instead, we establish the structure of these behaviours and provide a foundation for evaluating their functional relevance following release and, ultimately, for determining whether behavioural relationships expressed in captivity correspond to those favored in the wild.

**Table 1.**
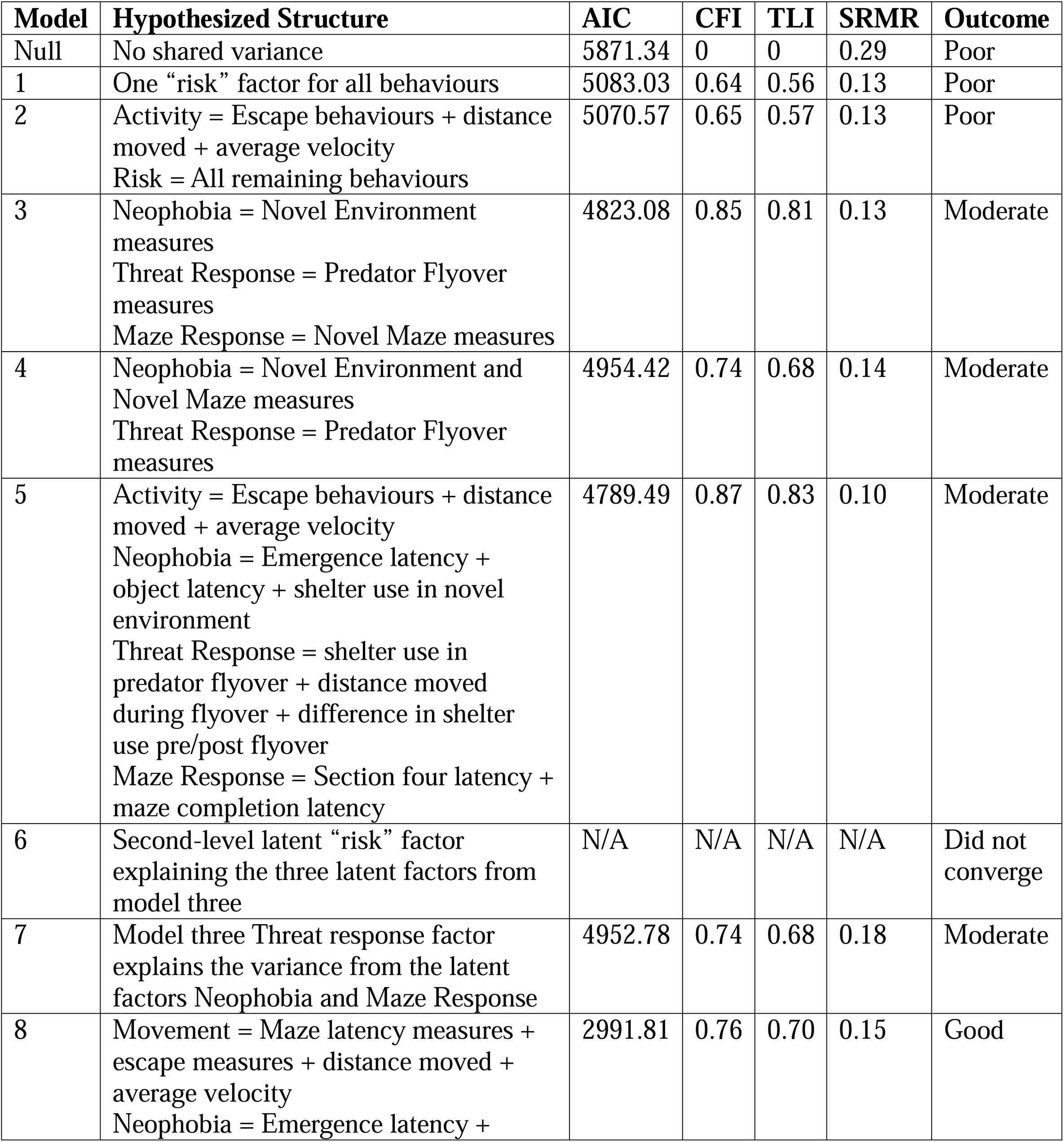
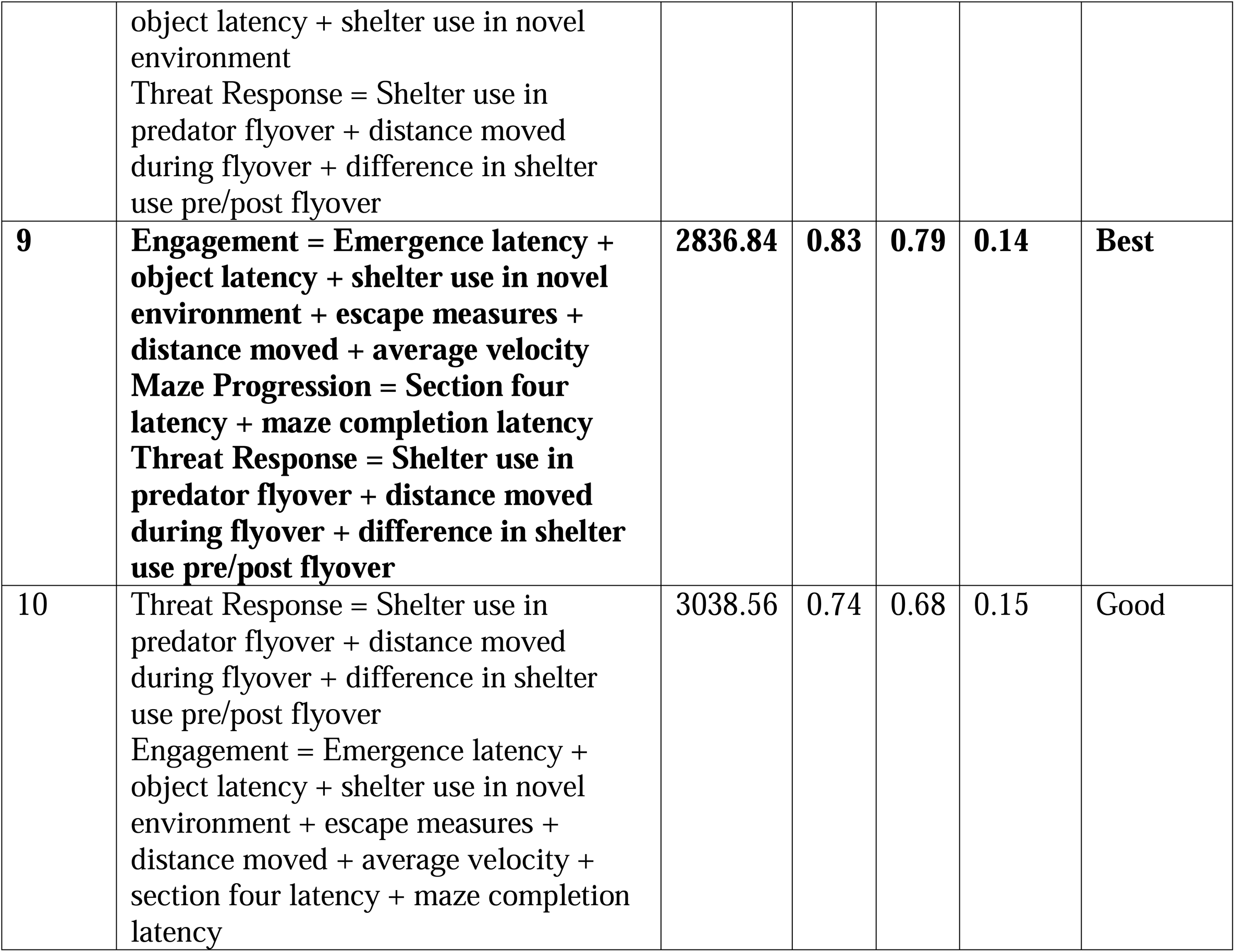
Comparison of candidate behavioural structure models ranging from a single general behavioural axis to assay-specific and functionally grouped latent factors. Model fit was evaluated using Akaike information criterion (AIC), comparative fit index (CFI), Tucker-Lewis index (TLI), and standardized root mean square residual (SRMR). Greater support is indicated by lower AIC values, higher CFI/TLI values, and lower SRMR values. The best-supported model was identified based on relative model performance and biological interpretability and is bolded.

| Model | Hypothesized Structure | AIC | CFI | TLI | SRMR | Outcome |
|---|---|---|---|---|---|---|
| Null | No shared variance | 5871.34 | 0 | 0 | 0.29 | Poor |
| 1 | One “risk” factor for all behaviours | 5083.03 | 0.64 | 0.56 | 0.13 | Poor |
| 2 | Activity = Escape behaviours + distance moved + average velocity<br>Risk = All remaining behaviours | 5070.57 | 0.65 | 0.57 | 0.13 | Poor |
| 3 | Neophobia = Novel Environment measures<br>Threat Response = Predator Flyover measures<br>Maze Response = Novel Maze measures | 4823.08 | 0.85 | 0.81 | 0.13 | Moderate |
| 4 | Neophobia = Novel Environment and Novel Maze measures<br>Threat Response = Predator Flyover measures | 4954.42 | 0.74 | 0.68 | 0.14 | Moderate |
| 5 | Activity = Escape behaviours + distance moved + average velocity<br>Neophobia = Emergence latency + object latency + shelter use in novel environment<br>Threat Response = shelter use in predator flyover + distance moved during flyover + difference in shelter use pre/post flyover<br>Maze Response = Section four latency + maze completion latency | 4789.49 | 0.87 | 0.83 | 0.10 | Moderate |
| 6 | Second-level latent “risk” factor explaining the three latent factors from model three | N/A | N/A | N/A | N/A | Did not converge |
| 7 | Model three Threat response factor explains the variance from the latent factors Neophobia and Maze Response | 4952.78 | 0.74 | 0.68 | 0.18 | Moderate |
| 8 | Movement = Maze latency measures + escape measures + distance moved + average velocity<br>Neophobia = Emergence latency + object latency + shelter use in novel environment<br>Threat Response = Shelter use in predator flyover + distance moved during flyover + difference in shelter use pre/post flyover | 2991.81 | 0.76 | 0.70 | 0.15 | Good |
| **9** | **Engagement = Emergence latency + object latency + shelter use in novel environment + escape measures + distance moved + average velocity<br>Maze Progression = Section four latency + maze completion latency<br>Threat Response = Shelter use in predator flyover + distance moved during flyover + difference in shelter use pre/post flyover** | **2836.84** | **0.83** | **0.79** | **0.14** | **Best** |
| 10 | Threat Response = Shelter use in predator flyover + distance moved during flyover + difference in shelter use pre/post flyover<br>Engagement = Emergence latency + object latency + shelter use in novel environment + escape measures + distance moved + average velocity + section four latency + maze completion latency | 3038.56 | 0.74 | 0.68 | 0.15 | Good |

## Methods

### Study Species

Over three breeding seasons (April – May, 2023 – 2025) we studied the behaviour of 257 juvenile APC hatched at the Houston Zoo’s captive breeding facility located at the NASA Johnson Space Center (Houston, Texas, USA). Captive rearing began in 1992 with 19 surviving founders, contributing to the four current captive breeding facilities that produce an estimated 300 birds annually to supplement the wild population (26,27). Although the number of captive breeders fluctuates annually, there are approximately 42 males and 54 females.

### Behavioural Trials

At two and four weeks of age, the behaviours (electronic supplementary material, Table S1) of individual APC chicks were quantified in a three-day testing battery that consisted of three independent assays: a novel environment and novel object (day one), a simulated predator flyover (day two), and a novel maze (day three). We videoed all ten-minute trials for behavioural transcription (N=1,332). Trials were run in the mornings, with some exceptions due to inclement weather (N=106). We recorded the temperature, windspeed, weather (six terms including Sunny, Cloudy, etc.), and humidity (only for 2024 and 2025) as described by the Apple iPhone weather app at the start of each trial. Whenever possible testing order of individuals was randomized but the architecture of some rearing cages necessitated haphazard chick capture. All behavioural assays took place in topless, modified brooder cages (81 cm x 97 cm x 31 cm; GQF Manufacturing 0534 - Universal Box Brooder) without the heating elements in place (electronic supplementary material, Appendix S1).

### Novel Environment and Object Assay

To ensure novelty, we covered the brooder sidewalls with fabric (2023) or coloured cardboard (2024 and 2025), using pink for trial one and blue for trial two. We placed one pink (trial one) or blue (trial two) novel object in the front half of the enclosure and another in the back half (ping pong balls in 2023 and rubber ducks in 2024 and 2025). The flooring had a 7 x 6 grid of non-slip shelf liner fabric. We covered each open brooder top with mesh to prevent escape. Each chick was placed in a wooden acclimation chamber (30 cm x 30 cm x 30 cm) attached to the outside of the testing arena for at least one minute before remotely opening a door that allowed it to access the novel environment. Chicks were captured immediately before testing and transported back to their home cages immediately following the trials.

### Predator Flyover Assay

Using the same brooder as the novel environment assay so that the arena was no longer novel, we replaced the novel objects with small trays of food (APC starter, Mazuri Exotic Animal Nutrition, St. Louis, MO, USA) to measure willingness to forage. Each bird was placed by hand in the top right corner of the arena at the start of the trial (electronic supplementary material, Appendix S1) and had access to the acclimation chamber for shelter through an open door. Four minutes into the trial, we “flew” a soaring silhouette of an aerial predator above the testing arena so birds could visually detect the moving silhouette and the associated changes in light as it crossed the arena. The silhouette was passed from right to left for trial one (red-tailed hawk; *Buteo jamaicensis*) and from left to right for trial two (great horned owl; *Bubo virginianus*) for an average duration of 7.5 seconds.

### Novel Maze Assay

We inserted four wooden panels to divide the brooder enclosure into a maze with five equal sections that the chick could access through panel openings at alternating ends of the sections. We placed each bird in the front right corner of the arena. At times, the maze walls blocked the camera’s view of the chick, but the distance moved and average velocity were interpreted from assumed linear movement of the positions before and after the chick was obscured. To prevent learning, we reversed the panel openings for the second trial.

### Behavioural Transcription

We used a mixture of fully manual transcription, manual movement tracking software (MTrackJ; ((28)), and machine learning open-source programs (DeepLabCuts (29,30) and SIMBA (31)). MTrackJ (version 1.5.1) encodes coordinates each time the user clicks on the subject, allowing for location and movement evaluation. DeepLabCuts (version 3.0.0rc7) automates the movement tracking of individuals using body position points. We used the open-source SIMBA program (version 4.2.8) to identify bouts of escape attempts but were unable to identify bouts of novel object interactions, so we manually recorded those behaviours. We achieved acceptable precision, recall, and F1 scores for all assays and visually checked outputs. See electronic supplementary material, Appendix S2 for details regarding parameters, model training, and evaluation.

### Statistical Analyses

We visually inspected continuous variables for substantial skew and applied log or square-root transformations where needed to improve conformity with Gaussian error assumptions. We added a 0.001 constant to numerical variables before transforming to eliminate zeroes. Latency measures with maximum scores for unobserved behaviours were treated as right-censored responses rather than assigning the maximum latency score to the video length (600 seconds). Transformations and response distributions for each measure are provided in electronic supplementary material, Table S1. After excluding individuals that successfully escaped the arena, the dataset contained 255 individuals.

We first modelled the variation of each behavioural measurement separately as a function of fixed effects (trial, year, deviation from average start time, weather, temperature, wind speed, deviation from average handling time, brooder ID, lighting on video footage (High, Medium, Low), presence of a glare (Y/N), whether the full enclosure was always in view (Y/N), and whether the camera shakes (Y/N) using the lme4 package (32) in RStudio (version 2025.09). These analyses quantified potential sources of measurement, environmental, and procedural variation, and we included a random intercept for rearing group ID to account for non-independence among individuals raised within the same group. We then calculated the adjusted repeatabilities (V_I/V_I + V_R, where V_I represents among-individual variance and V_R represents residual within-individual variance) for each variable and included the important fixed effects that influenced behavioural variation using the rptR package (33). Any behavioural measures that had zero among-individual variation were dropped from further analyses.

We fit all remaining behavioural measures into a multivariate mixed-effect model using the MCMCglmm package (34). For the latency variables, we fit a “cengaussian” distribution to account for individuals receiving a max latency score if they never engaged in the behaviour. We fit the rest of the variables with a “gaussian” distribution. We included the important fixed effects identified from the univariate models to account for measured nuisance conditions associated with individual behavioural measures in the model, as well as trial number to account for within-individual variation, so that the resulting covariance structure represented variation beyond the observed sources of extraneous variation. We fit the fixed effects at the trait level to ensure that no fixed effects from one assay were tested against variables of another assay and excluded observation-level fixed effects that significantly affected specific traits in the univariate models if their inclusion caused linear dependence in the multivariate matrix (rank deficiency). We retained only trial and year as fixed effects for all assays, and response-specific covariates were included where supported by the univariate analyses. See electronic supplementary material, Table S6, for explanations on which predictors were dropped from the multivariate model due to rank deficiency. We included random intercepts for individual identity and, after removing individuals with missing data for the fixed predictors, we ran the model on 248 individuals. Due to the complexity of our 13-trait multivariate model, we used a weakly informative, but regularizing covariance prior that shrunk the relative scale of among-individual variation (G) versus residual (R) variance but did not provide a well-defined mean (V_G_ = 0.05, V_R_=0.95, nu =14). We evaluated model robustness using three alternative weak priors that varied the relative G:R scale and degrees of freedom. We also assessed sensitivity to predictor exclusion by running the same model without the trait-level covariates identified from the univariate models (but including trial and year) as well as trait-level covariates derived from AIC model comparison. See supplementary material, Appendix S3 for additional information on the sensitivity analyses. Finally, we examined whether any measures taken across multiple assays (e.g., time spent attempting escape, latency to attempt escape, distance moved, average velocity) were repeatable across contexts through an additional multivariate model.

We extracted the posterior samples of the among-individual covariance G matrix from the multivariate mixed model. We fit each of the *a priori* models with confirmatory factor analyses (CFA), a specialized structural equation modeling technique for testing how well measured variables represent hypothesized latent factors, using the lavaan (35) and sem (36) packages, and compared fits using AIC-values. For the models grouping neophobic responses with movement and escape-related behaviours, we used the term “Engagement” to reflect how readily individuals initiated and sustained responses without implying a particular motivational state. “Threat Response” comprised predator-response specific behaviours and “Maze Progression” comprised the latencies to advance through the maze. We included one model that inferred the Threat Response factor as the overarching “risk” dimension, resulting in a model that proposed alternative hierarchical relationships (electronic supplementary material, Fig. S3).

To estimate among-individual variation for the best-supported latent factors while accounting for uncertainty, we extracted the individual-level posterior distributions from the covariance matrix (11). Using the latent factor loadings from the best performing model as the basis for factor extraction, we combined the posterior samples of the random effects with the factor loadings at each multivariate model iteration for each individual, resulting in a full posterior distribution of latent scores for each individual and factor, rather than a single point estimate. Note that for visualization purposes, we summarized the posterior distributions for each individual and trait by calculating the posterior mean, standard deviation, and 95% confidence interval. To assess within-individual variation between trials, or behavioural plasticity, we calculated the change in the behavioural measures with the strongest loadings by subtracting the trial one measured value from the trial two measured value for the individuals with two measurements (N=186) and performed a one-sample t-test on the difference. To determine whether the mean behavioural scores per latent factor differed by year, we ran a linear model with the mean score as the response variable and year as the predictor. To test whether the score variance differed by year, we ran an additional model that changed the variance assumption to unequal variance per year using the gls function from the nlme package (37,38) and compared model fit to a model of equal variance. We extracted the variance estimates to obtain the direction and magnitude of the behavioural distributions.

## Results

### Behavioural Repeatability and Predictors

There were 13 non-zero repeatable behavioural measures: five from the Novel Environment assay (emergence latency, latency to interact with a novel object, time spent in shelter, time spent attempting escape, and distance moved), six from the Predator Flyover assay (time spent attempting escape, average velocity, time spent in shelter, distance travelled during flyover, and the difference between time in shelter before versus after flyover), and three from the Novel Maze assay (latency to attempt escape, latency to enter section four, and latency to complete the maze). The adjusted repeatabilites, which measure the proportion of variance after controlling for fixed effects, for the 13 behaviours included in the multivariate model ranged from 0.095 to 0.371 (Table 2; electronic supplementary material, Fig. S4). Handling time, trial start time, weather, and wind speed explained variation in several behavioural measures, whereas technical video conditions affected relatively few retained measures. Full coefficients and confidence intervals are reported in electronic supplementary material, Tables S3 – S5.

**Table 2.** Repeatability estimates including 95% confidence intervals and covariates of behavioural measures included in the multivariate analysis.

| Assay | Behavioural Measure | Adjusted R (95% CI) | Covariates included in MCMCglmm |
|---|---|---|---|
| Novel Environment | Emergence Latency | 0.371 (0.252, 0.476) | Year, trial, weather, handling |
| Novel Environment | Object Latency | 0.216 (0.115, 0.341) | Year, trial |
| Novel Environment | Shelter Use | 0.23 (0.073, 0.356) | Year, trial, handling |
| Novel Environment | Time Attempting Escape | 0.233 (0.131, 0.354) | Year, trial |
| Novel Environment | Distance Moved | 0.095 (0.014, 0.266) | Year, trial, handling |
| Predator Flyover | Time Attempting Escape | 0.159 (0.014, 0.266) | Year, trial, start time |
| Predator Flyover | Average Velocity | 0.16 (0.026, 0.269) | Year, trial |
| Predator Flyover | Shelter Use | 0.357 (0.249, 0.466) | Year, trial, weather |
| Predator Flyover | Distance Moved during Flyover | 0.209 (0.09, 0.328) | Year, trial, handling, weather |
| Predator Flyover | Difference in Shelter Use Pre/Post Flyover | 0.347 (0.225, 0.463) | Year, trial |
| Novel Maze | Escape Attempt Latency | 0.168 (0.029, 0.294) | Year, trial, wind speed, weather |
| Novel Maze | Section Four Latency | 0.12 (0.026, 0.238) | Year, trial, start time, weather |
| Novel Maze | Latency to Complete Maze | 0.141 (0.009, 0.269) | Year, trial, camera movement, testing arena ID |

Assay type significantly predicted all measures in the univariate models. When tested in a separate multivariate model, the among-individual correlations between the same behavioural measures across different assays were generally weak, with posterior means near zero and credible intervals (39) overlapping zero, indicating limited cross-context consistency (electronic supplementary material, Table S2).

### Model Sensitivity

Our prior sensitivity analyses demonstrated that varying the G versus R variance affected repeatability estimates, which indicates that the variance partitioning is partially regularized by the prior. However, the posterior factor-score rankings were highly stable across all priors (Spearman rho ≥ 0.976 for our data-driven weak prior compared to our base prior and ≥ 0.890 under fully neutral priors), and the top-25 overlap remained high. The effective sample sizes were large for key variance terms, making it less likely that the results were MCMC artifacts.

The same three-factor structure was supported when models included only trial and year or covariates selected using AIC (electronic supplementary material, Appendix S3). Loading retained the same directions, and posterior scores were strongly correlated among approaches (Pearson *r* = 0.961 – 0.991), indicating that the inferred structure was robust to covariate specification. Including fixed effects ensured that the covariance structure represented variation beyond identified sources of extraneous variation.

### Models of Behavioural Structure

None of the models surpassed preferred model fit indices for comparative fit indices (CFI ≥ 0.90), Tucker-Lewis indices (TLI ≥ 0.90), and standardized root mean squared residuals (SRMR ≤ 0.08). Model five contained “Heywood cases”, which indicated that the model was too flexible or complex for the covariance structure, with over- or underfitting between latent factors, or that some traits load weakly on the proposed factors. Model six failed to estimate entirely.

Overall, the null model performed the worst. Models one, two, four, seven, and ten were also poor fits (Table 1). Model three and eight performed slightly better, but model nine had the most support, with the lowest AIC score, highest CFI and TLI, and the lowest SRMR. Patterns in the among-individual covariance matrix were consistent with the lack of support for a single risk-taking factor, with stronger covariance among subsets of behavioural measures (electronic supplementary material, Fig. S5). Model nine hypothesized three structural components: a) behavioural measures of the novel environment assay and the general activity measures (escape latency in the maze trials and time spent attempting escape in the predator trials) are best explained by a latent “Engagement” factor, b) that the measures during the predator flyover as well as the total time spent in the shelter are best explained by a latent “Threat Response” factor, and c) that the latency to enter section four of the maze and the latency to complete the maze are best explained by a latent “Maze Progression” factor (Fig. 1).

**Figure 1.**
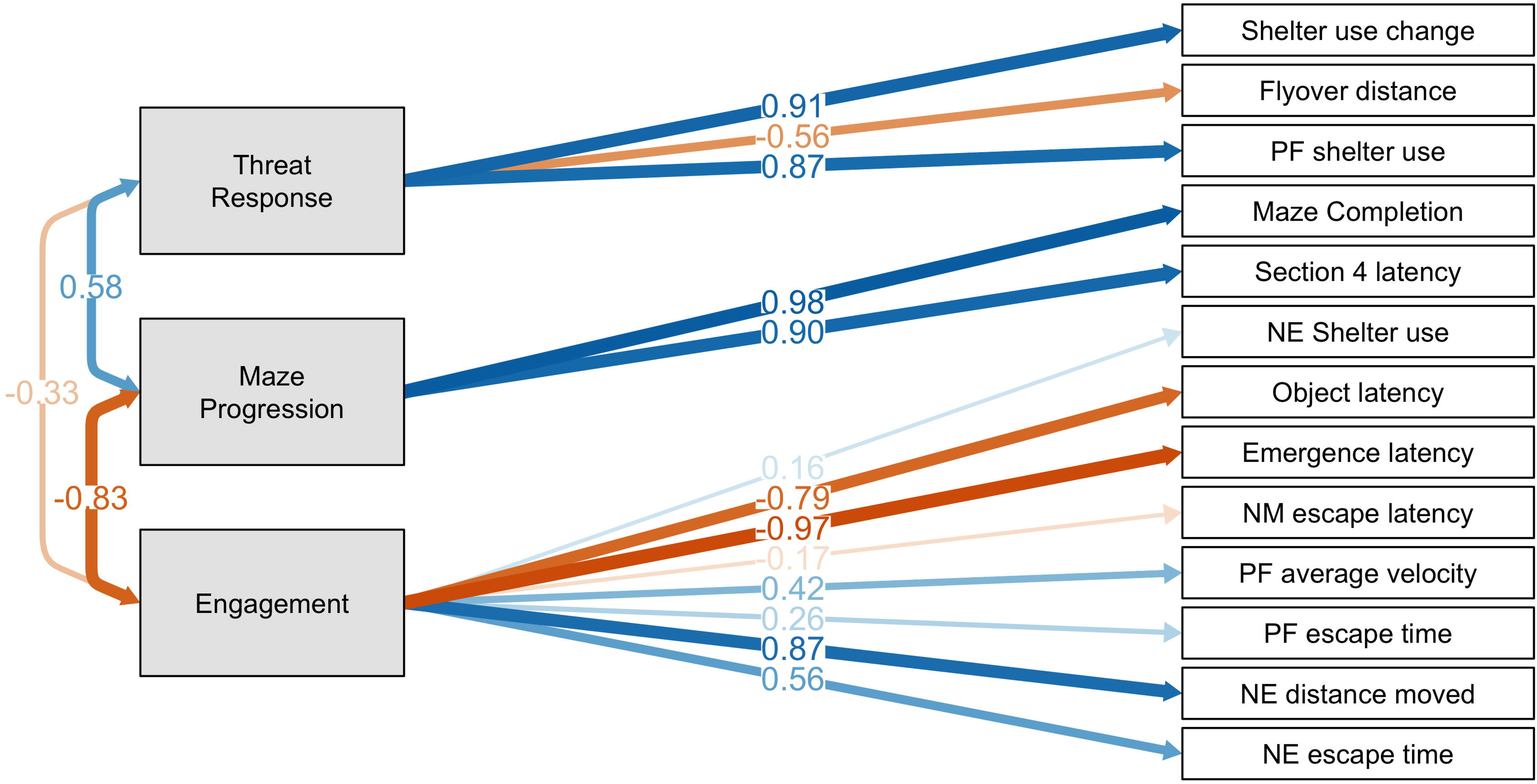
Structural equation model depicting relationships among latent and observed variables. Novel Environment and Object is abbreviated as NE, Predator Flyover is abbreviated as PF, and Novel Maze is abbreviated as NM. The large grey rectangles represent the three latent constructs, and white squares represent observed variables. Single-headed arrows indicate directional effects with path coefficients, and double-headed arrows represent covariances. The transparency of the arrows represents the strength of the directional effects and covariances, with blue indicating positive and orange indicating negative. Values correspond to standardized posterior mean loadings with 95% confidence intervals and are derived from the Bayesian MCMCglmm model. Model fit statistics: [CFI = 0.83, RLI = 0.79, SRMR = 0.14].

The Engagement factor had the most variable standardized loadings (-0.97 to 0.87), followed by Threat Response (-0.57 to 0.91), and Maze Progression (0.90 to 0.98; Table 3; electronic supplementary material, Fig. S6). After summarizing the among-individual variation across draws, the variation of all three factors (Engagement = 1.67, Threat Response = 1.05, and Maze Progression = 1.19) exceeded the summarized within-individual posterior uncertainty (Engagement = 1.21, Threat Response = 0.69, and Maze Progression = 0.87), indicating that the differences among individuals are distinguishable from estimation error.

**Table 3.** Behavioural measure loadings onto each latent factor from the best-supported model identified from confirmatory factor analysis. Year effect with its corresponding P value is included for each of the three latent factors. The estimated Pearson correlation coefficient is included to demonstrate the relationship between the latent factors, with arrows to indicate the direction of correlation.

| Latent Factor | Behavioural Measures | Standardized Loadings | Year Effect (P Value) | Relationships (Pearson Coefficient) |
|---|---|---|---|---|
| Engagement | Emergence latency | -0.97 | 0.076 | ↓ Threat Response (*r* = -0.44) |
|  | Distance Moved | 0.87 |  |  |
|  | Object Latency | -0.79 |  |  |
|  | Time attempting escape in novel environment | 0.56 |  |  |
|  | Average velocity in predator flyover | 0.42 |  |  |
|  | Time attempting escape in predator flyover | 0.25 |  |  |
|  | Latency to attempt escape in novel maze | -0.16 |  |  |
|  | Shelter use in novel environment | 0.16 |  |  |
| Threat Response | Difference in shelter use pre/post flyover | 0.91 | < 0.001 | ↑ Maze Progression (*r* = 0.78) |
|  | Shelter use during flyover trials | 0.88 |  |  |
|  | Distance moved during flyover | -0.573 |  |  |
| Maze Progression | Latency to complete maze | 0.98 | 0.011 | ↓ Engagement |
|  | Section four latency | 0.90 |  | (*r* = -0.88) |

### Behavioural Plasticity and Yearly Variation

Emergence latency decreased in trial two (*p* = 0.02, χ = -0.16), shelter use before versus after predator increased in trial two (*p* = 0.04, χ = 0.16), and the latency to enter section four of the maze increased in trial two (*p* = 0.04, χ = 0.19). Yearly cohorts also differed in the variation of their latent factor scores (Fig. 2). There was moderate support for variation in Engagement scores (*p* = 0.076) and significant support for Threat Response and Maze Progression scores (*p* < 0.001 and *p* = 0.0113, respectively). Score variation was highest for all latent factors in 2024 and lowest in 2023. Mean Engagement score decreased from 2023 to 2025, while mean Threat Response and Maze Progression scores increased from 2023 to 2025.

**Figure 2.**
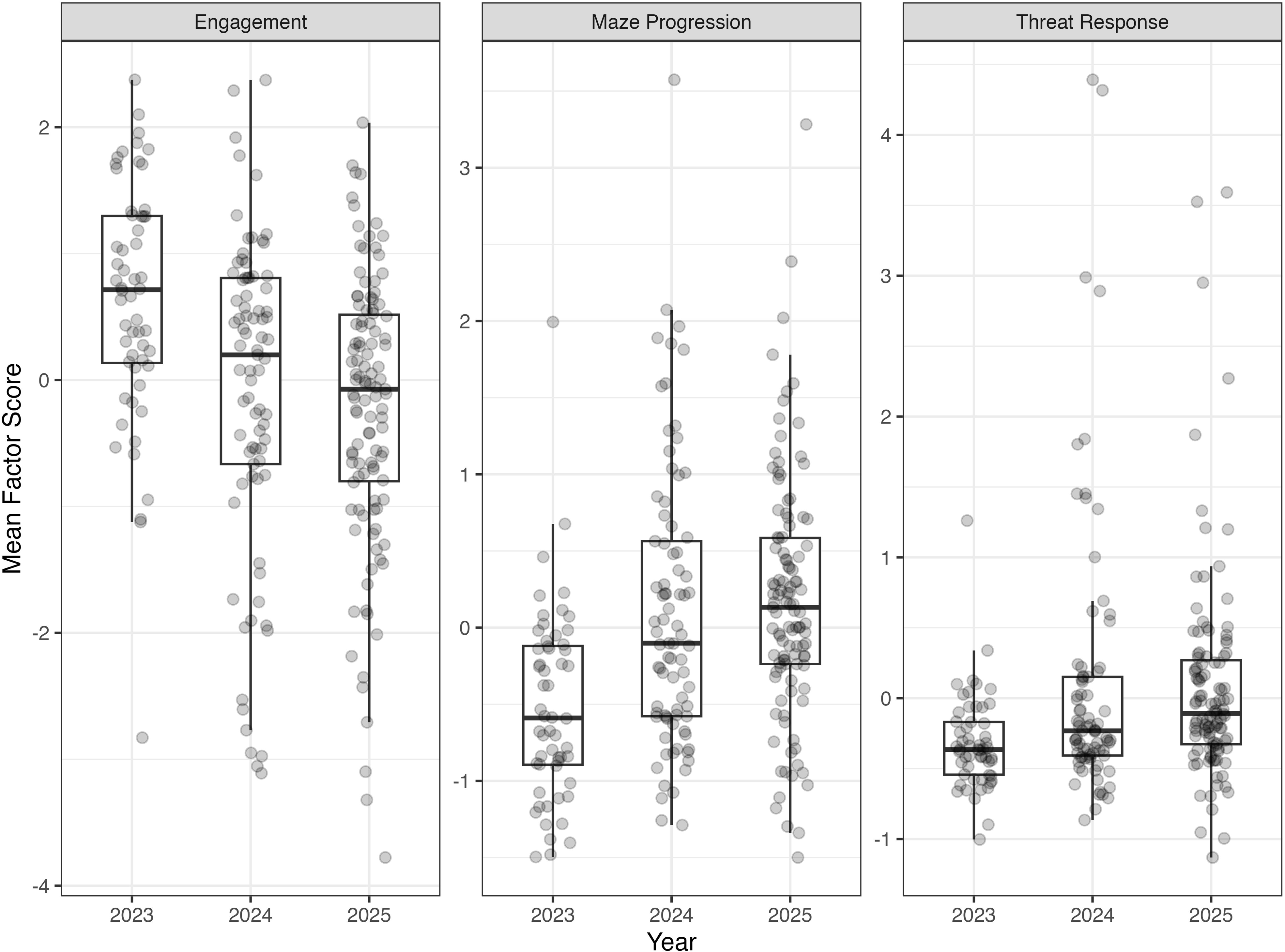
Boxplots show the distribution of individual mean posterior factor scores for the three latent behavioural axes *(a)* Engagement, *(b)* Threat Response, *(c)* Maze Progression. Points represent individual birds, with higher dispersion indicating greater among-individual behavioural variation within a cohort. Differences in score distributions were evaluated using generalized least squares models allowing year-specific variance structures.

### Latent Factor Relationships

To compare the relationship between the three latent factors, we fit unconstrained (different effects) of regressions into the factor models (Fig. 3, Table 3). As the latent factors were estimated from behavioural measures modeled on a transformed scale when appropriate, the relationships reported represent standardized associations among the latent behavioural dimensions rather than changes in the original units. Threat Response negatively predicted Engagement in ∼85% of the 574 posterior draws (mean std.all = -0.334), and positively predicted Maze Progression in ∼97% (mean std.all = 0.442). Maze Progression positively predicted Engagement in ∼97% (mean std.all = 0.989), and positively predicted Threat Response in ∼91% (mean std.all = 0.918). Engagement negatively predicted Threat Response in ∼81% of the draws (mean std.all = -0.413), and positively predicted Maze Progression in ∼97% (mean std.all = 0.67).

**Figure 3.**
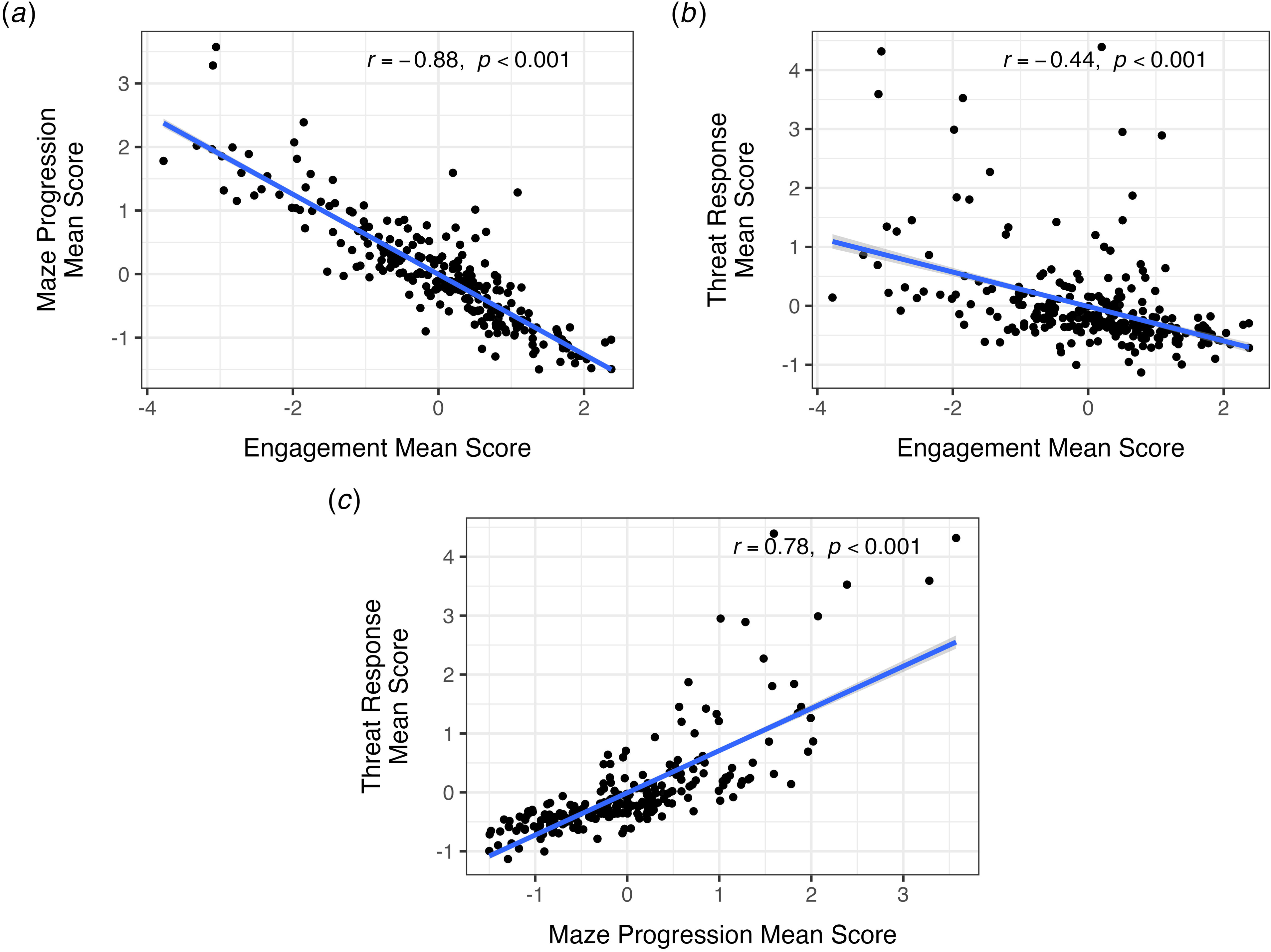
Pairwise relationships (on transformed units) between summed individual-level latent scores for Engagement, Maze Progression, and Threat response. Each point represents the mean posterior latent score for an individual estimated from the Bayesian MCMCglmm model. Chicks that emerge from the shelter more quickly (higher engagement scores) *(a)* enter maze sections sooner (lower maze progression scores), and *(b)* use the shelter less (lower threat response scores) during the predator flyover trials. *(c)* Chicks that enter the maze later (high maze progression scores) use the shelter more (higher threat response scores).

## Discussion

Our work addressed the central assumption underlying behavioural syndrome research that responses to novelty, predation risk, and exploration reflect a shared axis of risk-taking behaviour commonly referred to as boldness (40). By explicitly comparing alternative hypotheses of behavioural structure, we were able to show stronger support for a multidimensional organization of behaviour than for a single generalized risk-taking axis within a captive-bred population of Attwater’s prairie chickens. Behavioural responses were best explained by three partially independent latent dimensions representing engagement, threat response, and maze progression that were robust to manipulations of priors and covariates. Our findings suggest that risk-related behaviours in this system are more modular than commonly assumed, supporting growing evidence that behavioural syndromes may not represent fixed organismal properties, but instead context-dependent covariance structures shaped by ecological and selective environments (5,13,41,42).

The weak covariance observed among Engagement, Threat Response, and exploratory behaviours is particularly informative as these measures are frequently treated as interchangeable indicators of generalized boldness (but see (43,44)). Dimensionality-reduction approaches such as principal component analysis can summarize covariance among assays into a smaller number of statistical axes; however, interpreting that these axes reflect biologically unified behavioural syndromes requires an additional assumption that shared covariance reflects a common underlying behavioural process (11). However, the lack of strong integration among these latent dimensions within our study system suggests that responses to novelty and predation risk may instead arise from partially distinct underlying processes. This distinction may help explain why studies examining the fitness consequences of boldness often report inconsistent or contradictory results across systems (3,4). If the covariance structure among behaviours differs among populations or environments, then behavioural assays that appear to measure the same generalized trait may in fact capture different underlying axes of variation. However, although model nine was best supported, it did not exceed all conventional fit thresholds despite extensive model exploration. Therefore, these dimensions should not be interpreted as discrete or exhaustive behavioural types, but rather as the best-supported representation of covariance among the behavioural measures considered. The lack of an ideal fit may itself reflect the biological complexity underlying risk-related behaviours, where responses are unlikely to conform perfectly to simplified categorical frameworks such as “boldness”. Loading magnitudes also varied with covariate manipulation, underscoring the value of controlling procedural variation, although factor direction and overall structure remained robust.

Interestingly, although several individual behaviours were not themselves repeatable across assays (e.g., escape-related measures), they loaded consistently onto the same latent Engagement dimension. This pattern suggests that individuals may express broader behavioural tendencies through different context-specific behaviours rather than through rigidly fixed responses. Such patterns may reflect mechanisms similar to trait compensation, where individuals achieve similar functional outcomes through alternative behavioural pathways (13). This further complicates attempts to define generalized behavioural types using single assays and highlights the importance of evaluating covariance structure directly rather than assuming that superficially similar assays measure the same trait.

Although predation pressure and environmental unpredictability may favor the integration of behavioural responses across contexts in the wild, reinforcing covariance among antipredator behaviour, exploration, and novelty responses (45), whether the supported multidimensional structure arose in captivity cannot be determined without comparable wild APC data. Nevertheless, captivity alters predation risk, resource predictability, social environments, and human exposure, each of which could affect the strength of behavioural covariance (16,24,25,46). The trend in our average latent factor scores from 2023 – 2025 may suggest that behavioural expression shifts across captive generations or management conditions; however, additional work is needed to determine whether these patterns reflect directional responses to captivity or broader changes in behavioural structure. Rather than simply producing “bolder” or “shyer” individuals, captivity may reshape the covariance matrix underlying behavioural syndromes, effectively “unbundling” functionally linked behaviours. Our repeatability estimates further support this dynamic interpretation. The average repeatability estimates were lower than the 0.35 average estimated across behavioural studies (47), while suggestive evidence of within-individual plasticity in key traits implies that individuals retained substantial flexibility in their responses across trials. Together, these findings align with reaction norm perspectives emphasizing that behavioural expression reflects both consistent among-individual differences and environmentally responsive plasticity (10,48).

These results have important implications for conservation breeding and translocation programs as captive and wild phenotypes are often assumed to be comparable (25,46). Captive breeding initiatives typically prioritize preserving genetic diversity yet dedicate relatively little attention to preserving behavioural structure. If behavioural covariance structures can be modified under captive conditions, then behavioural phenotypes expressed in captivity may result in undesired responses to wild selection pressures. This could have compounded effects for population growth if the underlying genetic structure does not produce the expected behavioural correlation and magnitudes when captive individuals breed with wild counterparts, as seen when crossing captive mice lineages (7). Additionally, when environmental conditions differ substantially from those experienced under natural selection, behavioural plasticity itself may become an important component of translocation success by allowing reintroduced individuals to respond optimally to novel post-release conditions. To determine the role of plasticity in translocations, and the effects of captive genetic structure on population growth, further work examining the consistency of behavioural types across several repeated assays (>2) and the behaviours of the population post-release are necessary.

The possibility of capitalizing on behavioural types for species recovery, even without considering behavioural structure, is not unfounded. Translocation studies on swift foxes (*Vulpes velox;* (17)), Tasmanian devils (*Sarcophilus harrissii;* (16)), brushtail possums (*Trichosurus vulpecula;* (19)), desert tortoises (*Gopherus agassizii;* (20)), and greater bilbies (*Macrotis lagotis;* (18)) have found significant links between behavioural variation and key variables for recovery success, including survival and reproduction. However, different aspects of boldness were found to be predictors of success; for example, neophobic behaviours were related to breeding success in greater bilbies (18), but unrelated to survival, whereas neophobic responses in desert tortoises did predict survival (20). Our findings suggest that any behavioural screening programs may benefit from moving beyond generalized classifications such as “bold” or “shy”. Because different components of risk-related behaviours may be semi-independent, decisions based on a single assay or measure may fail to capture the behavioural traits most relevant for survival after release. Instead, behavioural assessments may be most informative when tailored to the primary ecological challenges faced by individuals at release sites. For example, when predation represents the dominant source of mortality, as in the APC, antipredator responses may be more relevant than exploratory tendencies or novelty response behaviours, whereas dispersal, social integration, or resource acquisition may represent stronger selective pressures in other systems. Identifying how these behavioural dimensions covary, whether they remain stable across captive and wild environments, how they translate from parent to offspring, and how they are linked to fitness will allow managers to more effectively incorporate behavioural variation into conservation decision-making.

## Conclusion

Our results highlight the importance of explicitly testing assumptions about behavioural structure before applying behavioural syndrome frameworks (11). Our study suggests that covariance among risk-related behaviours in captive APC were better represented by three partially independent dimensions rather than by a single boldness axis. This result cautions treating responses to novelty, predators, and exploration as interchangeable measures, particularly when behavioural assessments inform translocation decisions. Conservation translocation programs provide a valuable opportunity to not only evaluate the ecological consequences of behavioural variation, but also to understand whether altered selective environments reshape behavioural architecture, which may be as important as understanding shifts in mean behavioural expression for improving conservation outcomes. Future work identifying the mechanisms that maintain, disrupt, or restructure behavioural covariance will be essential for understanding how behavioural syndromes evolve and how behavioural diversity can be effectively incorporated into conservation management.

## Acknowledgements

We are grateful to Brenton Laing for his instruction in the use of machine learning methods to collect behavioural data and to Ned Dochtermann and the SQuID working group for statistical advice. This work would not have been possible without the practical and administrative support of the Houston Zoo and its animal care staff led by Kyle McAuliffe as the recovery program coordinator, the APC recovery team with Dr. Jim Muller acting as the Recovery Team Leader, and the U.S. Fish and Wildlife Service staff. This is publication 48 of the Center for Biodiversity & Conservation Research at the University of Mississippi.

