## Supplemental figures and appendices for "Deconstructing “Boldness”: Latent behavioural structure across novelty, predation, and exploration in an endangered species translocation program"

*Proceedings B,* doi:10.1098/rspb

### Electronic Supplementary Material

Contents **Page**

**Appendix S1:** Representative behavioural arena images. 2

**Appendix S2:** Behavioural Transcription using Machine Learning Methods 5

**Appendix S3:** Sensitivity Analysis of Multivariate Model 7

**Table S1:** Ethogram of measures collected and the corresponding behavioural assay. 9

**Table S2:** Summary of the multivariate model comparing the behavioural consistency of measures collected across multiple assays. 11

**Table S3:** Estimated sources of variation with 95% confidence intervals (CI) from univariate fixed effect models and the adjusted repeatability for the five Novel Environment and Object variables. 12

**Table S4:** Estimated sources of variation with 95% confidence intervals (CI) from univariate fixed effect models and the adjusted repeatability for the six Predator Flyover variables. 15

**Table S5:** Estimated sources of variation with 95% confidence intervals (CI) from univariate fixed effect models and the adjusted repeatability for the three Novel Maze variables 18

**Table S6:** Explanation of fixed effects significant univariate models but dropped in multivariate analysis. 20

**Figures S1-S3:** Eleven *a priori* models (hypotheses) to explain the covariance structure among behaviours commonly associated with "boldness" including neophobic, anti-predator avoidance, and exploration. 21

**Figure S4:** Univariate repeatability estimates (R) of the 13 behavioural measures included in the multivariate analysis. 26

**Figure S5:** Covariance heatmap of behavioural measures included in the multivariate analysis to demonstrate the potential groupings of traits in the proposed models. 27

**Figure S6:** Magnitude (0-1) and direction (-/+) of behavioural measure loading onto the three latent factors identified from the best supported *a priori* model. 28

**Supplementary References** 29

**Appendix S1.** Behavioural trial diagrams

We have included labeled photos of our behavioural assay arenas for each trial. The left (or top) pink panel is trial 1 from 2023 and the right (or bottom) blue panel is trial 2 from 2025.

Novel Environment Assay. We placed individual chicks in the acclimation chamber as indicated in the following figure. The novel objects were placed in the front and back of the arena on the right and left. The red circle on the right panel indicates the light source for the arena. There is no circle on the left panel as the light source was above the camera frame.

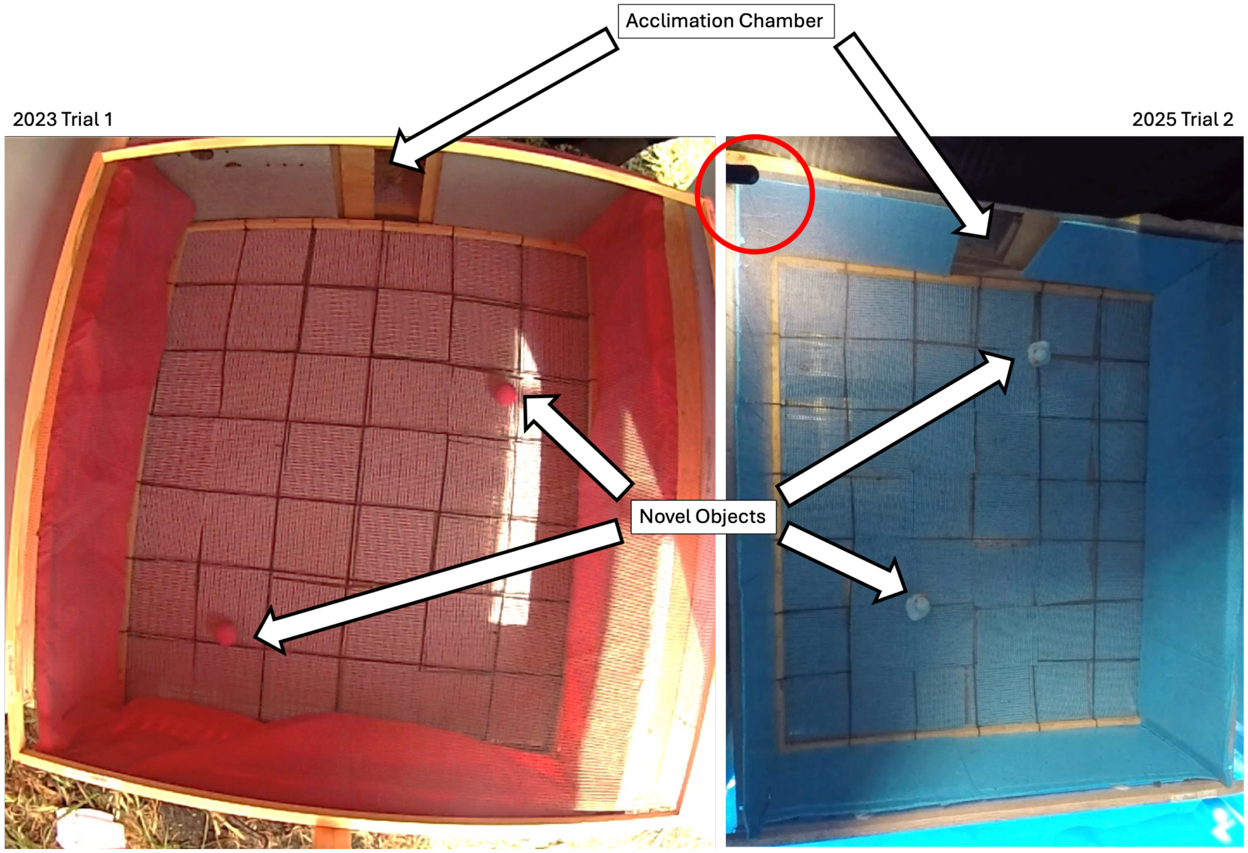

Predator Flyover. The doors to the acclimation chamber remained open for shelter use, but we placed individual chicks in the area indicated by the green circles on the right and left panels. The food trays were either rectangle or circular but mimicked the food trays that were included in their home enclosures. The red circle on the right panel again indicates the light source for the arena.

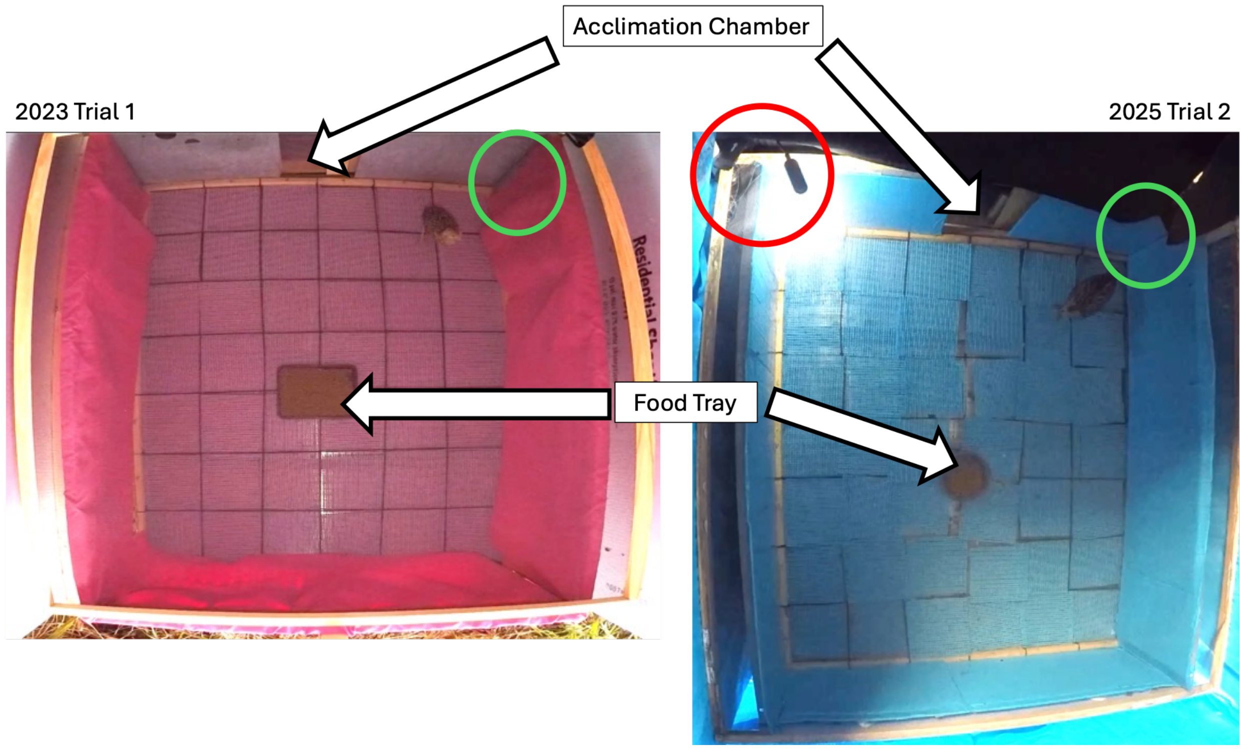

Novel Maze. The acclimation chamber doors remained closed for this assay to encourage exploration. We placed individual chicks in the area indicated by the green circles at the start of the trial. The grid flooring was removed to fit the wooden dividers, and we altered the divider openings to create a maze. We reversed the opening order for trial two as shown in the bottom panel.

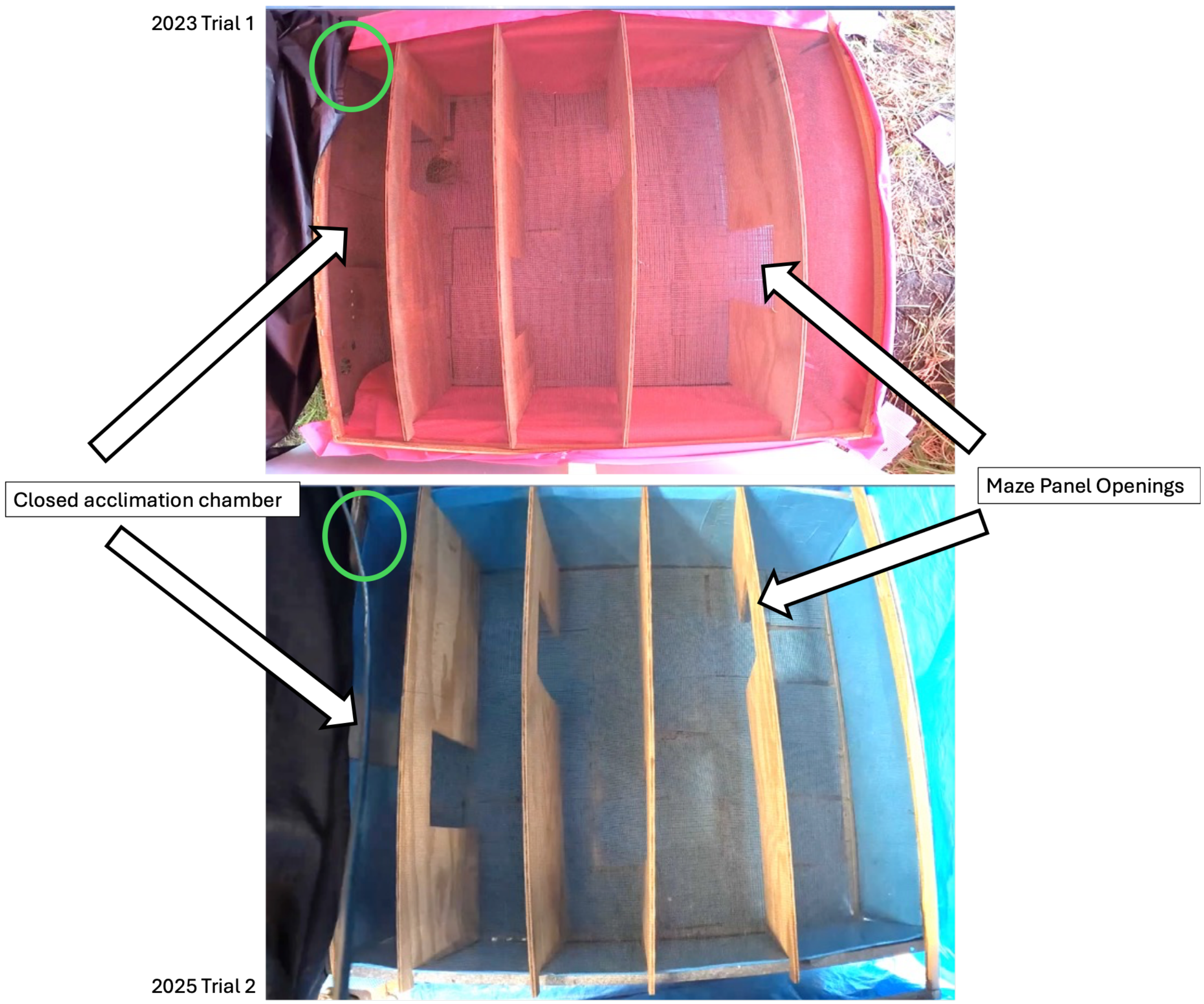

**Appendix S2.** Behavioural Transcription using Machine Learning Methods

Given that each assay consisted of differing levels of bird visibility, we used assay-specific video subsets for machine learning methods. We labeled frames from 27 mixed-quality Novel Environment videos from 2023 and 2024 (N=271), 48 mixed-quality Predator Flyover (N=270), and 48 mixed-quality Novel Maze (N=267). The remaining 632 videos from 2025 were not used in the training dataset since training occurred prior to 2025 (211 Novel Environment videos, 211 Predator Flyover videos, and 210 Novel Maze videos). We used 95% of the labeled training dataset to train the machine learning model using the PyTorch engine in DeepLabCuts and the remaining 5% to test its performance. We began with 20 training videos per assay and increased the training dataset based on visual inspection of tracking performance. We did not utilize outlier correction due to the number of frames with missing subjects. For the Novel Environment videos, we achieved a 0.74 precision, 0.69 recall, and 0.72 F1 score with the following parameters: 2000 random forest estimators, sqrt max features, gini criterion, 0.2 test size, bout split type, 5 minimum sample leaf, and a balanced class weight. With the same parameters, we achieved a 0.71 precision, 0.7 recall, and a 0.71 F1 score for the Predator Flyover videos. We used the same parameters except for class balance for the Novel Maze videos, which performed better with a class weight of 10 for Present and a class weight of 1 for Absent (Precision = 0.73, Recall = 0.54, F1 = 0.62). We inspected detection accuracy with manually and semi-manually transcribed videos and performed Kleinberg smoothing filtering on the Predator Flyover and Novel Maze videos to improve detection.

The best performing neural network for pose estimation was the DlcrnetStride16Ms5 (1) with default parameters for all assays. We obtained an average root mean square error (RMSE) with a 0.6 P cutoff of 4.44 for the training dataset and 7.94 for the test dataset for the Novel Environment videos and a RMSE with 0.6 P cutoff of 2.29 for the training videos and 7.81 for the test videos for the Predator Flyover videos. Given that numerically evaluating an acceptable RMSE depends on several factors (2), it is best to visually check the tracking output of the videos to ensure accuracy, which we did for all videos. The Novel Maze videos improved visually when ran with the Predator Flyover training model rather than the maze trained model.

For behavioral extraction of the 1,440 videos from 257 individuals, the SIMBA model incorrectly identified 274 (19%) videos as either having escape bouts when there were none or not having escape bouts when bouts were present. It identified 46 (3.2%) videos where we manually missed escape bouts that were present. All incorrect identifications and missed bouts were corrected in the final dataset. In addition to the extracted variables, we also tracked instances in the footage where there were levels of low lighting, glares, or when part of the enclosure walls were out of frame. Across all 1,438 videos, 2.0% had low lighting, 7.5% contained a glare, and 50.0% had at least a portion of the upper enclosure outside the camera's field of view, primarily affecting visibility during escape attempts. In instances such as the maze assay where the chick might be obscured by the maze panel, the distance measurements were estimated from the last known position as a linear distance. Consequently, these estimates may underestimate actual distance traveled if chicks followed a non-linear path while obscured.

**Appendix S3.** Sensitivity Analysis of Multivariate Model

We assessed the sensitivity to weak inverse-Wishart priors that varied the degrees of freedom and the relative scale of G versus R variance. The priors we tested included a unit scale weak prior since the traits were standardized ($V_G = 1$, $V_R=1$, nu =14), a slightly more regularizing weak prior with a finite mean ($V_G = 0.05$, $V_R=0.95$, nu =18), and a data-driven weak prior based on observed variance ($V_G = 0.25$, $V_R=0.75$, nu =14).

Due to competing statistical approaches for the use of univariate model analysis of fixed effects and their resulting inclusion in the larger multivariate model, we performed two sensitivity analyses to ensure that our biological conclusions remained robust. First, we repeated our multivariate analysis using an a priori model excluding the screened nuisance predictors and retained only trial and year as we considered those to be biologically relevant to our question. After comparing the 11 *a priori* models of latent structure with the updated covariance matrix, we found that the same latent model was supported and all latent-variable loadings retained the same direction, although the magnitudes did differ.

We also elected to run another sensitivity analysis using an information theoretic approach of the univariate models (3). We first ran a complete model with all predictors for each of the 13 behaviors included in the multivariate analysis, checked multicollinearity, and dropped variables with high VIF estimates. We then refit the reduced model, rechecked collinearity, and ran the dredge function in the MuMIn package (4) with trial and year as fixed effects to compare the large number of alternative models. We used model averaging for the top models that were within two AICs of the best model and compared the relative importance of each variable. We then included those predictors in the final model and reran our multivariate analysis with those predictors derived from information theory rather than significance tests. The same latent factor model remained the most supported with the covariance matrix updated to include AIC-derived fixed effects. The posterior latent scores from the original covariate-selection approach using univariate models were highly correlated with both the no-fixed-effect sensitivity model and the new information-theoretic approach ($r$ = 0.961 – 0.991). The AIC-derived scores correlated with the original scores at r = 0.980 – 0.989, with Spearman correlations of 0.949 – 0.979. The posterior SDs were also highly concordant ($r$ = 0.941 – 0.957).

**Table S1.** Ethogram of measures collected and the corresponding behavioural assay. Behaviours with non-zero repeatability that were included in multivariate analysis are bolded along with the associated assay. Continuous responses were standardized to a mean of zero and a standard deviation of one after any transformations. "Censored Gaussian" refers to right-censored latency responses in the multivariate MCMCglmm analysis; all other retained responses were modeled with Gaussian families.

| **Operational Description** | **What it measures** | **Assay Observed** | **Analysis Treatment** |
|---|---|---|---|
| **Emergence latency** | Time before chick exits the acclimation chamber from when the door opens. Censored at 600 seconds, or the length of the video | **Novel Environment & Object** | **Log; censored Gaussian** |
| **Object latency** | Time until chick approaches the novel object in the Novel Environment trials. Censored at 600 seconds, or the length of the video | **Novel Environment & Object** | **Square root; censored Gaussian** |
| **Shelter time** | The time spent within the acclimation chamber during the Novel Environment and Predator Flyover trials | **Novel Environment & Object**<br>**Predator Flyover** | **NE: Log; Gaussian**<br>**PF: Log, Gaussian** |
| **Escape latency** | Time before attempting to jump out of the enclosure from the point of the door opening in the Novel Environment trials and the bird placement in the arena in the Maze and Predator Flyover trials. Censored at 600 seconds, or the length of the video | Novel Environment & Object<br>Predator Flyover<br>**Novel Maze** | NE: Log<br>PF: Log<br>**NM: untransformed, censored Gaussian** |
| **Time Escape** | The time spent attempting to jump out of the enclosure | **Novel Environment & Object**<br>**Predator Flyover**<br>Novel Maze | NE: untransformed, Gaussian<br>PF: Square-root, Gaussian<br>NM: Square-root |
| **Distance moved** | Total distance moved during the trial (in the form of a path from movement tracking) | **Novel Environment & Object**<br>Predator Flyover<br>Novel Maze | **NE: log, Gaussian**<br>PF: Log<br>NM: untransformed |

| | | | |
|---|---|---|---|
| **Average velocity** | The average distance per pixel across the trials | Novel Environment & Object<br>**Predator Flyover**<br>Novel Maze | NE: log;<br>PF: Log, Gaussian<br>**NM: Log** |
| Difference in distance moved | The difference in the distance moved during the last minute of the trial compared to the first minute of the trial | Predator Flyover | Untransformed |
| Difference in average velocity | The difference in the average distance per pixel during the last minute of the trial compared to the first minute of the trial | Predator Flyover | Untransformed |
| Time spent in food tray | Time spent in the food tray region of interest during the Predator Flyover trials | Predator Flyover | Log |
| Flyover escape | Time spent attempting to jump out of the enclosure during the predator flyover event | Predator Flyover | Untransformed |
| Flyover food | Time spent in the food tray region of interest during the predator flyover event | Predator Flyover | Log |
| Flyover shelter | Time spent in the acclimation chamber during the predator flyover event | Predator Flyover | Untransformed |
| **Flyover distance** | Distance moved during the predator flyover event | **Predator Flyover** | **Log, Gaussian** |
| Flyover velocity | Average velocity during the predator flyover event | Predator Flyover | Log |
| **Difference in shelter use** | Difference in the time spent in the acclimation chamber after the predator flyover event compared to before the predator flyover event | **Predator Flyover** | **Untransformed; Gaussian** |
| Section two latency | The time from the placement of the bird in the maze until it enters the second section of the maze. Censored at 600 | Novel Maze | Log |

| | | | |
|---|---|---|---|
| | seconds, or the length of the video | | |
| Section three latency | The time from the placement of the bird in the maze until it enters the third section of the maze. Censored at 600 seconds, or the length of the video | Novel Maze | Log |
| **Section four latency** | The time from the placement of the bird in the maze until it enters the fourth section of the maze. Censored at 600 seconds, or the length of the video | **Novel Maze** | **Untransformed; censored Gaussian** |
| **Latency to complete maze** | The time from the placement of the bird in the maze until it enters the fifth (final) section of the maze, thus completing it. Censored at 600 seconds, or the length of the video | **Novel Maze** | **Untransformed; censored Gaussian** |

**Table S2.** Summary of the multivariate model comparing the behavioural consistency of measures collected across multiple assays. Measures are presented in pairwise comparisons with the mean, 95% confidence interval, and associated probability that the correlation is greater than zero. Estimates are derived from extracting the posterior correlation matrices, storing the posterior correlations as an array and summarizing the posterior correlations for each pair.

| **Trait 1** | **Trait 2** | **Mean** | **95% CI** | **P (r>0)** |
|---|---|---|---|---|
| NE Escape Latency | PF Escape Latency | -0.258 | -0.933, 0.836 | 0.312 |
| NE Escape Latency | NM Escape Latency | -0.377 | -0.948, 0.718 | 0.241 |
| PF Escape Latency | NM Escape Latency | 0.250 | -0.694, 0.907 | 0.693 |
| NE Escape Time | PF Escape Time | 0.051 | -0.453, 0.554 | 0.574 |
| NE Escape Time | NM Escape Time | 0.056 | -0.448, 0.561 | 0.579 |
| PF Escape Time | NM Escape Time | 1 | 1 | 1 |
| NE Shelter Time | PF Shelter Time | 0.322 | -0.178, 0.706 | 0.909 |

**Table S3.** Estimated sources of variation with 95% confidence intervals (CI) from univariate fixed effect models and the adjusted repeatability for the five Novel Environment and Object variables. Random intercepts were included for individual chick and rearing group and the associated p-value for each variable is reported. The fixed effects included are trial 2 (1-2), year (2023, 2024, 2025), weather (isolated thunderstorm, partially sunny, sunny, windy), brooder ID (A, B, C), deviation from average handling time, glare presence (Y/N), footage lighting (high, medium, low), enclosure entirely in camera frame (Y/N), and shorter than ten minute video (Y/N).

| | **Emergence Latency** | | **Object Latency** | | **Shelter Time** | | **Escape Time** | | **Distance Moved** | |
|---|---|---|---|---|---|---|---|---|---|---|
| **Fixed Effects** | **Estimate (95% CI)** | **P value** | **Estimate (95% CI)** | **P value** | **Estimate (95% CI)** | **P value** | **Estimate (95% CI)** | **P value** | **Estimate (95% CI)** | **P value** |
| **Intercept** | -231.73 (-682.4, 171.36) | 0.310 | -490.04 (-846.12, -139.61) | 0.009 | -98.99 (-476.15, 271.40) | 0.616 | 331.54 (-71.27, 731.14) | 0.112 | 265.99 -119.01, 667.68) | 0.194 |
| **Trial: 2** | -0.06 (-0.20, 0.07) | 0.328 | 0.08 (-0.09, 0.24) | 0.352 | -0.20 (0.37, -0.03) | 0.024 | -0.13 (-0.30, 0.04) | 0.143 | -0.06 (-0.23, 0.12) | 0.534 |
| **Year** | 0.11 (-0.08, 0.34) | 0.310 | 0.24 (0.07, 0.42) | 0.009 | 0.05 (-0.13, 0.24) | 0.617 | -0.16 (-0.36, 0.04) | 0.112 | -0.13 (-0.33, 0.06) | 0.194 |
| **Weather: Isolated Thunderstorms** | 0.58 (0.08, 1.05) | 0.018 | 0.28 (-0.27, 0.81) | 0.312 | 0.05 (-0.53, 0.60) | 0.854 | -0.10 (-0.67, 0.44) | 0.734 | 0.18 (-0.42, 0.74) | 0.550 |
| **Weather: Partially Sunny** | -0.66 (-1.20, -0.14) | 0.015 | -0.38 (-0.88, 0.16) | 0.154 | -0.44 (-0.97, 0.14) | 0.114 | 0.34 (-0.19, 0.91) | 0.230 | 0.36 (-0.24, 0.89) | 0.214 |
| **Weather: Sunny** | -0.05 (-0.22, 0.13) | 0.591 | -0.12 (-0.32, 0.12) | 0.247 | -0.02 (-0.22, 0.21) | 0.837 | -0.18 (-0.38, 0.04) | 0.097 | 0.31 (0.10, 0.51) | 0.005 |
| **Weather: Windy** | -0.03 (-0.50, 0.44) | 0.915 | 0.08 (-0.43, 0.68) | 0.761 | -0.51 (-1.05, 0.11) | 0.076 | 0.05 (-0.50, 0.67) | 0.864 | 0.79 (0.23, 1.36) | 0.008 |

| | | | | | | | | | | |
|---|---|---|---|---|---|---|---|---|---|---|
| **Brooder_id: B** | -0.34 (-0.54, -0.13) | 0.001 | -0.19 (-0.41, 0.05) | 0.107 | 0.16 (-0.08, 0.41) | 0.191 | 0.09 (-0.15, 0.33) | 0.443 | 0.010 (-0.14, 0.34) | 0.417 |
| **Brooder_id: C** | -0.14 (-0.35, 0.05) | 0.162 | 0.02 (-0.20, 0.24) | 0.842 | 0.08 (-0.16, 0.31) | 0.505 | 0.11 (-0.13, 0.33) | 0.374 | -0.05 (-0.28, 0.19) | 0.666 |
| **Handling Deviation** | 0.18 (0.07, 0.29) | 0.001 | 0.09 (-0.01, 0.19) | 0.090 | -0.18 (-0.29, 0.07) | 0.001 | 0.02 (-0.09, 0.14) | 0.672 | -0.15 (-0.27, -0.04) | 0.008 |
| **Glare: N** | 0.39 (-0.73, 1.60) | 0.501 | -0.74 (-1.84, 0.44) | 0.196 | 0.25 (-0.92, 1.46) | 0.669 | 0.14 (-0.99, 1.31) | 0.814 | -0.13 (-1.28, 1.06) | 0.822 |
| **Glare: Y** | -0.03 (-1.17, 1.25) | 0.956 | -0.58 (-1.76, 0.63) | 0.336 | 0.16 (-1.08, 1.44) | 0.798 | 0.52 (-0.66, 1.78) | 0.396 | 0.04 (-1.20, 1.29) | 0.943 |
| **Lighting: High** | -0.05 (-0.43, 0.32) | 0.786 | -0.26 (-0.71, 0.16) | 0.226 | 0.18 (-0.30, 0.62) | 0.442 | -0.13 (-0.59, 0.30) | 0.553 | -0.17 (-0.61, 0.26) | 0.442 |
| **Lighting: Low** | -0.12 (-0.64, 0.43) | 0.654 | -0.12 (-0.70, 0.52) | 0.712 | 0.00 (-0.63, 0.70) | 0.996 | -0.45 (-1.05, 0.20) | 0.172 | -0.08 (-0.73, 0.60) | 0.821 |
| **Enclosure In View: N** | -0.05 (-0.29, 0.17) | 0.630 | 0.339 (0.10, 0.57) | 0.003 | 0.17 (-0.08, 0.40) | 0.17 | -0.13 (-0.37, 0.11) | 0.290 | -0.05 (-0.29, 0.20) | 0.662 |
| **Short Video: Y** | - | - | - | - | - | - | -0.38 (-0.79, 0.05) | 0.070 | - | - |
| **Random Effects** | **Std. Dev. (95% CI)** | **Variance** | **Std. Dev. (95% CI)** | **Variance** | **Std. Dev. (95% CI)** | **Variance** | **Std. Dev. (95% CI)** | **Variance** | **Std. Dev. (95% CI)** | **Variance** |
| **Individual** | 0.35 (0.18, 0.46) | 0.12 | 0.25 (0, 0.42) | 0.062 | 0.37 (0.14, 0.52) | 0.139 | 0.29 (0, 0.45) | 0.082 | 0.00 (0, 0) | 0.000 |
| **Rearing Group** | 0.48 (0.33, 0.62) | 0.23 | 0.27 (0.10, 0.39) | 0.075 | 0.30 (0.12, 0.42) | 0.087 | 0.34 (0.19, 0.47) | 0.119 | 0.35 (0.19, 0.47) | 0.121 |
| **Residual** | 0.68 (0.61, 0.74) | 0.46 | 0.85 (0.77, 0.93) | 0.729 | 0.86 (0.78, 0.95) | 0.738 | 0.85 (0.77, 0.94) | 0.729 | 0.92 (0.85, 0.98) | 0.832 |

| **Repeatability** | **R (95% CI)** | - | **R (95% CI)** | - | **R (95% CI)** | - | **R (95% CI)** | - | **R (95% CI)** | - |
|---|---|---|---|---|---|---|---|---|---|---|
| | 0.371 (0.252, 0.476) | - | 0.216 (0.115, 0.341) | - | 0.23 (0.073, 0.356) | - | 0.233 (0.131, 0.354) | - | 0.095 (0.014, 0.266) | - |

**Table S4.** Estimated sources of variation with 95% confidence intervals (CI) from univariate fixed effect models and the adjusted repeatability for the six Predator Flyover variables. Random intercepts were included for individual chick and rearing group and the associated p-value for each variable is reported. The fixed effects included are trial number (1-2), year (2023, 2024, 2025), weather (cloudy, isolated thunderstorm, partially sunny, rainy, windy), deviation from average start time, deviation from average handling time, lighting (low, medium, high), glare presence on footage (Y/N), enclosure entirely in camera view (Y/N), and presence of camera movement on footage (Y/N).

| | **Escape Time** | | **Average Velocity** | | **Shelter Time** | | **Flyover Distance Moved** | | **Difference Shelter Time** | |
|---|---|---|---|---|---|---|---|---|---|---|
| **Fixed Effects** | **Estimate (95% CI)** | **P value** | **Estimate (95% CI)** | **P value** | **Estimate (95% CI)** | **P value** | **Estimate (95% CI)** | **P value** | **Estimate (95% CI)** | **P value** |
| **Intercept** | -126.35 (-523.59, 267.56) | 0.544 | 257.66 (-86.90, 569.62) | 0.127 | -167.61 (-550.15, 179.83) | 0.368 | 258.43 (-89.00, 572.75) | 0.119 | -23.55 (-71.14, 19.21) | 0.312 |
| **Trial: 2** | -0.08 (-0.26, 0.09) | 0.379 | -0.32 (-0.49, -0.16) | 0.000 | -0.08 (-0.25, 0.08) | 0.373 | -0.01 (-0.20, 0.16) | 0.882 | 0.02 (-0, 0.04) | 0.069 |
| **Year** | 0.06 (-0.13, 0.26) | 0.544 | -0.13 (-0.28, 0.04) | 0.127 | 0.08 (-0.09, 0.27) | 0.368 | -0.13 (-0.28, 0.04) | 0.119 | 0.01 (-0.01, 0.04) | 0.312 |
| **Weather: Cloudy** | -0.06 (-0.28, 0.18) | 0.583 | 0.07 (-0.12, 0.28) | 0.458 | -0.17 (-0.37, 0.05) | 0.08 | 0.01 (-0.20, 0.23) | 0.918 | -0.01 (-0.03, 0.02) | 0.473 |
| **Weather: Isolated Thunderstorms** | 0.55 (-0.12, 1.16) | 0.095 | -0.43 (-1.01, 0.13) | 0.148 | 0.66 (0.08, 1.25) | 0.030 | -0.58 (-1.19, 0.03) | 0.058 | -0.03 (-0.10, 0.05) | 0.460 |
| **Weather: Partially Sunny** | 0.32 (-0.42, 1.09) | 0.413 | 0.28 (-0.39, 0.98) | 0.423 | 0.16 (-0.59, 0.90) | 0.669 | 0.06 (-0.66, 0.78) | 0.866 | 0.03 (-0.07, 0.12) | 0.580 |
| **Weather: Rainy** | -1.02 (-2.18, 0.21) | 0.088 | 0.56 | 0.307 | -0.38 (-1.51, 0.83) | 0.502 | 0.43 (-0.71, 1.60) | 0.452 | -0.06 (-0.21, 0.07) | 0.403 |

| | | | | | | | | | | |
|---|---|---|---|---|---|---|---|---|---|---|
| | | | (-0.49, 1.72) | | | | | | | |
| **Weather: Windy** | -0.43 (-1.61, 0.81) | 0.462 | -0.63 (-1.73, 0.50) | 0.248 | -0.49 (-1.59, 0.70) | 0.394 | 0.09 (-1.01, 1.27) | 0.874 | 0.02 (-0.11, 0.15) | 0.765 |
| **Start Time Deviation** | -0.00 (-0.01, -0.00) | 0.004 | -0.00 (-0.00, 0.00) | 0.697 | 0.00 (-0.00, 0.00) | 0.410 | 0.00 (-0.00, 0.00) | 0.996 | -0.00 (-0, 0) | 0.187 |
| **Handling Deviation** | 0.05 (-0.08, 0.19) | 0.442 | -0.07 (-0.19, 0.04) | 0.230 | 0.04 (-0.08, 0.17) | 0.487 | -0.15 (-0.27, -0.03) | 0.014 | 0.01 (-0.01, 0.02) | 0.364 |
| **Lighting: Low** | -1.26 (-2.68, 0.13) | 0.070 | 0.75 (-0.54, 2.07) | 0.262 | -0.88 (-2.21, 0.47) | 0.209 | -0.92 (-2.23, 0.49) | 0.191 | -0.03 (-0.20, 0.13) | 0.767 |
| **Lighting: Medium** | 0.23 (-0.12, 0.56) | 0.196 | 0.27 (-0.09, 0.59) | 0.106 | 0.04 (-0.31, 0.38) | 0.810 | 0.03 (-0.33, 0.35) | 0.874 | -0.00 (-0.04, 0.04) | 0.940 |
| **Glare: Y** | 0.43 (0.01, 0.86) | 0.035 | 0.38 (-0.02, 0.78) | 0.053 | -0.01 (-0.44, 0.41) | 0.970 | -0.13 (-0.55, 0.30) | 0.521 | -0.00 (-0.05, 0.05) | 0.918 |
| **Enclosure In View: N** | 0.15 (-0.07, 0.38) | 0.184 | 0.05 (-0.14, 0.26) | 0.599 | 0.24 (0.03, 0.45) | 0.028 | 0.05 (-0.15, 0.26) | 0.603 | 0.03 (0, 0.05) | 0.046 |
| **Camera Shakes : Y** | -0.15 (-1.09, 0.77) | 0.764 | -1.82 (-2.73, -0.93) | 0.00 | -0.28 (-1.23, 0.66) | 0.571 | -0.77 (-1.72, 0.18) | 0.115 | -0.00 (-0.12, 0.13) | 0.970 |
| **Random Effects** | **Std. Dev. (95% CI)** | **Variance** | **Std. Dev. (95% CI)** | **Variance** | **Std. Dev. (95% CI)** | **Variance** | **Std. Dev. (95% CI)** | **Variance** | **Std. Dev. (95% CI)** | **Variance** |
| **Individual** | 0.29 (0, 0.47) | 0.082 | 0.35 (0, 0.50) | 0.122 | 0.59 (0.42, 0.71) | 0.348 | 0.33 (0, 0.49) | 0.111 | 0.07 (0.04, 0.08) | 0.004 |
| **Rearing Group** | 0.31 (0.12, 0.44) | 0.094 | 0.11 (0, 0..27) | 0.013 | 0.08 (0, 0.28) | 0.007 | 0.00 (0, 0.22) | 0.000 | 0.02 (0, 0.04) | 0.000 |
| **Residual** | 0.89 (0.80, 0.97) | 0.788 | 0.85 | 0.715 | 0.82 (0.74, 0.90) | 0.670 | 0.90 (0.81, 0.98) | 0.810 | 0.10 (0.09, 0.11) | 0.010 |

| | | | | | | | | | |
|---|---|---|---|---|---|---|---|---|---|
| | | | (0.76, 0.93) | | | | | | |
| **Repeatability** | | | | | | | | | |
| | 0.159 (0.014, 0.266) | | 0.16 (0.026, 0.269) | | 0.357 (0.249, 0.466) | | 0.209 (0.09, 0.328) | | 0.347 (0.225, 0.463) |

**Table S5.** Estimated sources of variation with 95% confidence intervals (CI) from univariate fixed effect models and the adjusted repeatability for the three Novel Maze variables. Random intercepts were included for individual chick and rearing group and the associated p-value for each variable is reported. The fixed effects included are trial number (1-2), year (2023, 2024, 2025), deviation from average start time, weather (isolated thunderstorms, partially sunny, rainy, sunny, windy), wind speed, brooder ID (A, B, C), glare presence on footage (Y/N), lighting on footage (high, medium, low), presence of camera movement on footage (Y/N), and problem with arena setup visible on footage such as disturbed maze dividers (Y/N).

| | **Escape Latency** | | **Latency Section 4** | | **Latency Section 5** | |
|---|---|---|---|---|---|---|
| **Fixed Effects** | **Estimate (95% CI)** | **P value** | **Estimate (95% CI)** | **P value** | **Estimate (95% CI)** | **P value** |
| **Intercept** | -363.53 (-740.89, -12.77) | 0.064 | -750.58 (-1097.87, -400.47) | 0.000 | -721.14 (-1113.36, -369.31) | 0.000 |
| **Trial: 2** | 0.52 (0.29, 0.72) | 0.00 | 0.22 (0.01, 0.44) | 0.044 | 0.13 (-0.07, 0.33) | 0.242 |
| **Year** | 0.18 (0.01, 0.37) | 0.064 | 0.37 (0.20, 0.54) | 0.000 | 0.36 (0.18, 0.55) | 0.000 |
| **Start Time Deviation** | - | - | 0.53 (0.01, 1.01) | 0.034 | 0.00 (-0, 0) | 0.088 |
| **Weather: Isolated Thunderstorms** | 0.11 (-0.40, 0.60) | 0.659 | 0.42 (-0.13, 0.98) | 0.144 | 0.60 (0.14, 1.08) | 0.017 |
| **Weather: Partially Sunny** | -0.26 (-0.87, 0.35) | 0.411 | 0.27 (-0.31, 0.83) | 0.343 | 0.47 (-0.12, 1.08) | 0.116 |
| **Weather: Rainy** | 0.77 (0.019, 1.34) | 0.010 | -0.09 (-0.30, 0.13) | 0.450 | 0.20 (-0.33, 0.77) | 0.491 |
| **Weather: Sunny** | -0.07 (-0.29, 0.18) | 0.577 | 0.31 (-0.20, 0.79) | 0.218 | 0.02 (-0.19, 0.24) | 0.877 |
| **Weather: Windy** | 0.47 (-0.04, 1.00) | 0.068 | 0.00 (0, 0) | 0.059 | 0.40 (-0.12, 0.92) | 0.121 |
| **Wind Speed** | -0.04 (-0.08, -0.01) | 0.019 | -0.01 (-0.40, 0.02) | 0.570 | -0.02 (-0.06, 0.01) | 0.165 |
| **Brooder ID: B** | 0.05 (-0.17, 0.28) | 0.650 | 0.18 (-0.06, 0.40) | 0.127 | 0.22 (-0.01, 0.46) | 0.061 |

| | | | | | | |
|---|---|---|---|---|---|---|
| **Brooder ID: C** | -0.03 (-0.25, 0.22) | 0.829 | 0.02 (-0.21, 0.24) | 0.855 | -0.02 (-0.25, 0.22) | 0.887 |
| **Glare: Y** | 0.13 (-0.29, 0.57) | 0.551 | 0.25 (-0.16, 0.64) | 0.225 | 0.24 (-0.17, 0.66) | 0.255 |
| **Lighting: Low** | 0.19 (-0.35, 0.79) | 0.513 | 0.33 (-0.22, 0.95) | 0.258 | 0.34 (-0.21, 0.90) | 0.245 |
| **Lighting: Medium** | 0.08 (-0.19, 0.37) | 0.605 | 0.11 (-0.17, 0.41) | 0.429 | 0.10 (-0.19, 0.40) | 0.479 |
| **Enclosure In View: N** | -0.03 (-0.29, 0.21) | 0.835 | 0.16 (-0.08, 0.40) | 0.185 | 0.17 (-0.08, 0.42) | 0.185 |
| **Camera Shakes : Y** | 0.30 (-0.14, 0.74) | 0.165 | 0.51 (0.08, 0.91) | 0.016 | 0.43 (0.01, 0.85) | 0.041 |
| **Problem Setup: Y** | - | - | -0.26 (-0.65, 0.15) | 0.210 | 0.09 (-0.32, 0.49) | 0.686 |
| **Random Effects** | **Std. Dev. (95% CI)** | **Variance** | **Std. Dev. (95% CI)** | **Variance** | **Std. Dev. (95% CI)** | **Variance** |
| **Individual** | 0.23 (0, 0.40) | 0.051 | 0.22 (0, 0.41) | 0.048 | 0.26 (0, 0.44) | 0.068 |
| **Rearing Group** | 0.31 (0.14, 0.42) | 0.095 | 0.24 (0, 0.36) | 0.058 | 0.25 (0, 0.37) | 0.061 |
| **Residual** | 0.87 (0.79, 0.94) | 0.757 | 0.86 (0.78, 0.93) | 0.743 | 0.86 (0.77, 0.93) | 0.741 |
| **Repeatability** | | - | | | | |
| | 0.168 (0.029, 0.294) | | 0.12 (0.026, 0.238) | | 0.141 (0.009, 0.269) | |

**Table S6.** Explanation of fixed effects significant univariate models but dropped in multivariate analysis. Some variables were significant in the full rank univariate models, but non-estimable in the multivariate model due to over-parameterization and resulting rank deficiency. Additional estimable covariates were removed to reduce model complexity when inclusion resulted in lack of model convergence, poor mixing, or less reliable variance estimates. Novel Environment is represented as NE, Predator Flyover as PF, and Novel Maze as NM. Fixed effects retained in final multivariate model are bolded.

| Trait | Fixed Effect | Decision |
|---|---|---|
| NE Emergence Latency | Brooder ID | Removed to reduce model complexity |
| **NE Emergence Latency** | **Handling Deviation** | **Retained** |
| **NE Emergence Latency** | **Weather** | **Retained** |
| NE Object Latency | Enclosure In View | Removed to reduce model complexity |
| **NE Shelter Use** | **Handling Deviation** | **Retained** |
| **NE Distance Moved** | **Handling Deviation** | **Retained** |
| NE Distance Moved | Weather | Non-estimable due to rank deficiency |
| **PF Time Attempting Escape** | **Start Time Deviation** | **Retained** |
| PF Time Attempting Escape | Glare Presence | Removed to reduce model complexity |
| PF Average Velocity | Camera Shake | Removed to reduce model complexity |
| PF Shelter Use | Enclosure In View | Removed to reduce model complexity |
| **PF Shelter Use** | **Weather** | **Retained** |
| **PF Distance Moved during Flyover** | **Deviation Handling** | **Retained** |
| **PF Distance Moved during Flyover** | **Weather** | **Retained** |
| PF Difference in Shelter Use Pre/Post Flyover | Enclosure In View | Non-estimable due to rank deficiency |
| **NM Escape Attempt Latency** | **Weather** | **Retained** |
| **NM Escape Latency** | **Wind Speed** | **Retained** |
| **NM Section Four Latency** | **Weather** | **Retained** |
| **NM Section Four Latency** | **Start Time Deviation** | **Retained** |
| NM Section Four Latency | Camera Shake | Non-estimable due to rank deficiency |
| **NM Maze Completion Latency** | **Brooder ID** | **Retained** |
| **NM Maze Completion Latency** | **Camera Shake** | **Retained** |
| NM Maze Completion Latency | Weather | Non-estimable due to rank deficiency |

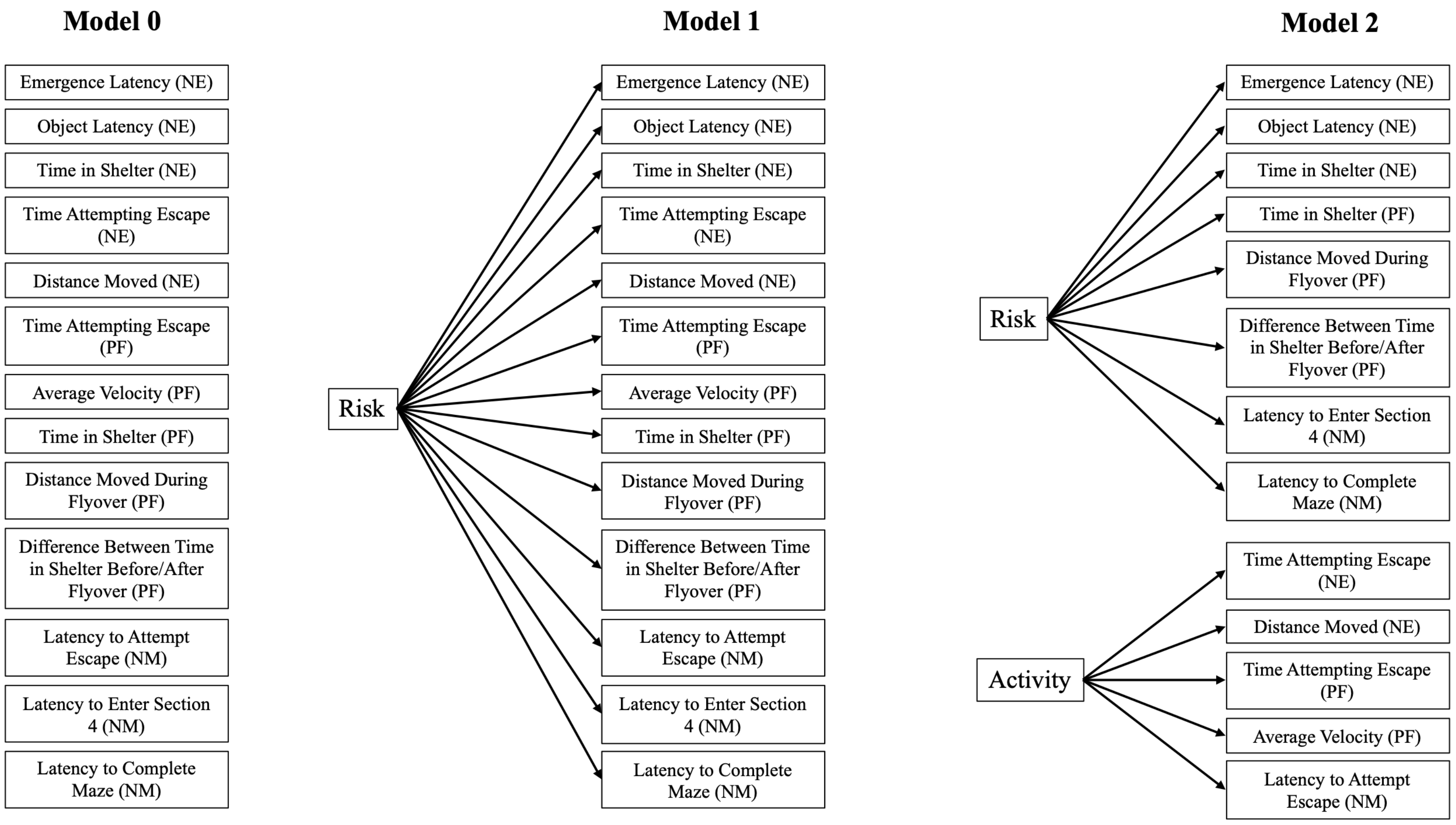

**Figure S1.** Models 0 – 2 of the eleven *a priori* models (hypotheses) to explain the covariance structure among behaviours commonly associated with "boldness" including neophobic, anti-predator avoidance, and exploration. Model zero is the null model assuming no shared variance; model one assumes one shared latent factor for all measures; and model two separates out activity behaviours from risk-related behaviours.

**Model 3**

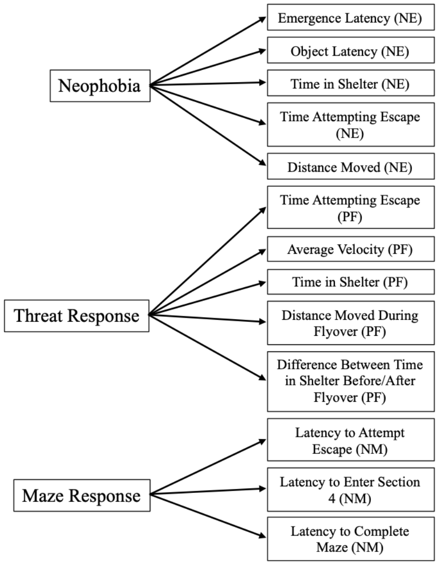

**Model 4**

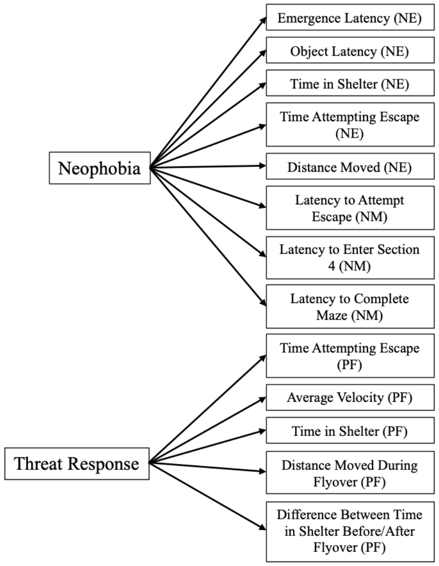

**Model 5**

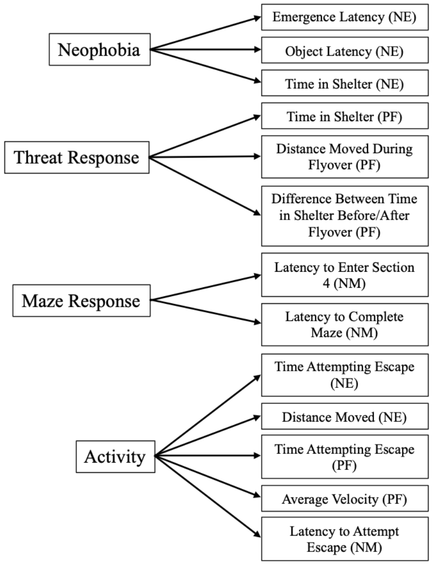

**Model 6**

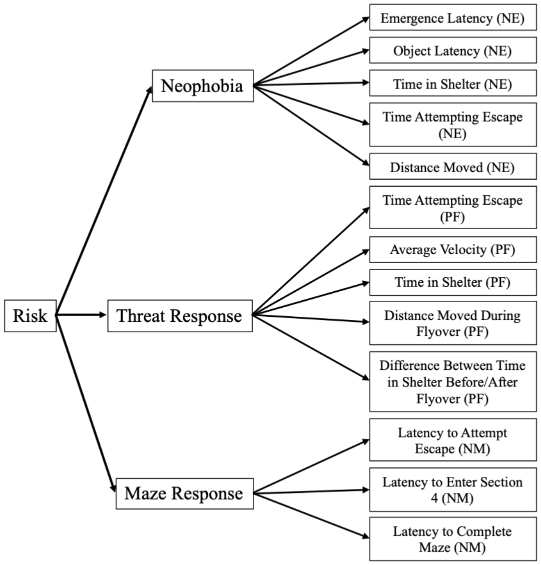

**Figure S2.** Models 3 – 6 of the eleven *a priori* models (hypotheses) to explain the covariance structure among behaviours commonly associated with "boldness" including neophobic, anti-predator avoidance, and exploration. Model three separates out behaviours by assay; model four combines neophobic and exploration behaviours but keeps predator avoidance measures separate; model five retains a general activity factor, but separates out the remaining measures by assay; and model six builds on model three through the addition of a second-level risk-factor explaining relationships between three latent factors.

**Model 7**

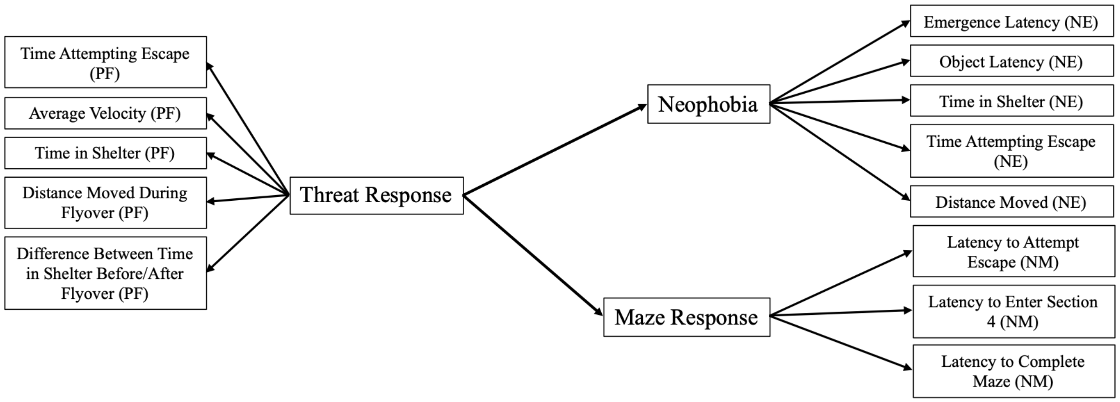

**Model 8**

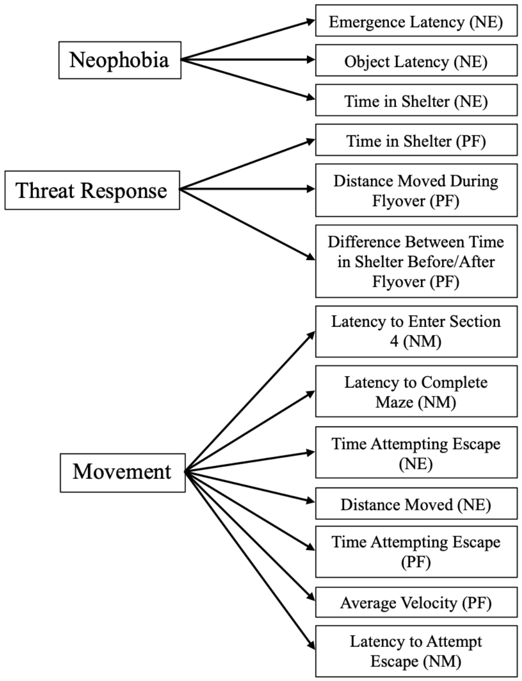

**Model 9**

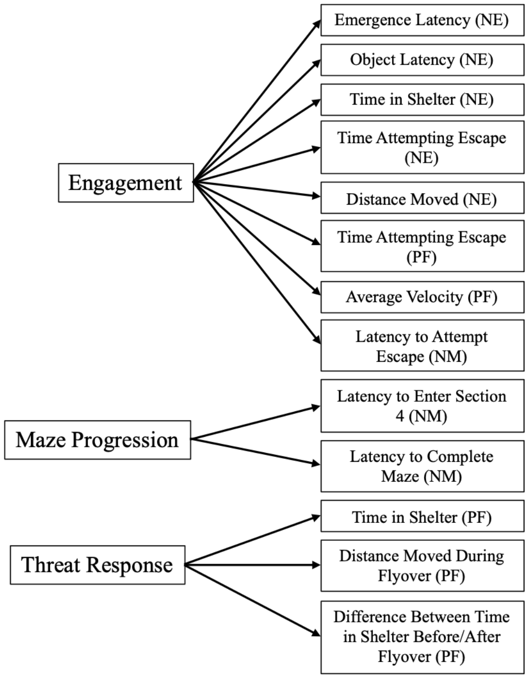

**Model 10**

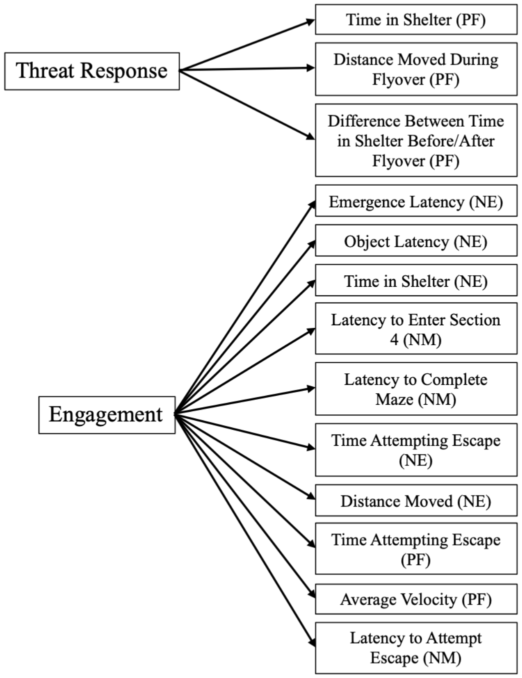

**Figure S3.** Models 7 – 10 of the eleven *a priori* models (hypotheses) to explain the covariance structure among behaviours commonly associated with "boldness" including neophobic, anti-predator avoidance, and exploration. Model seven proposes that predator avoidance measures explain the covariance structure between neophobic and exploratory measures; model eight combines maze and novel environment activity measures into "Movement", but keeps neophobic and predator avoidance specific measures separate; model nine proposes a combined activity and neophobic factor "Engagement", but keeps predator avoidance and maze exploration measures distinct; and model ten proposes a combination of "Engagement" and maze exploration measures into one factor, but keeps predator avoidance measures separate.

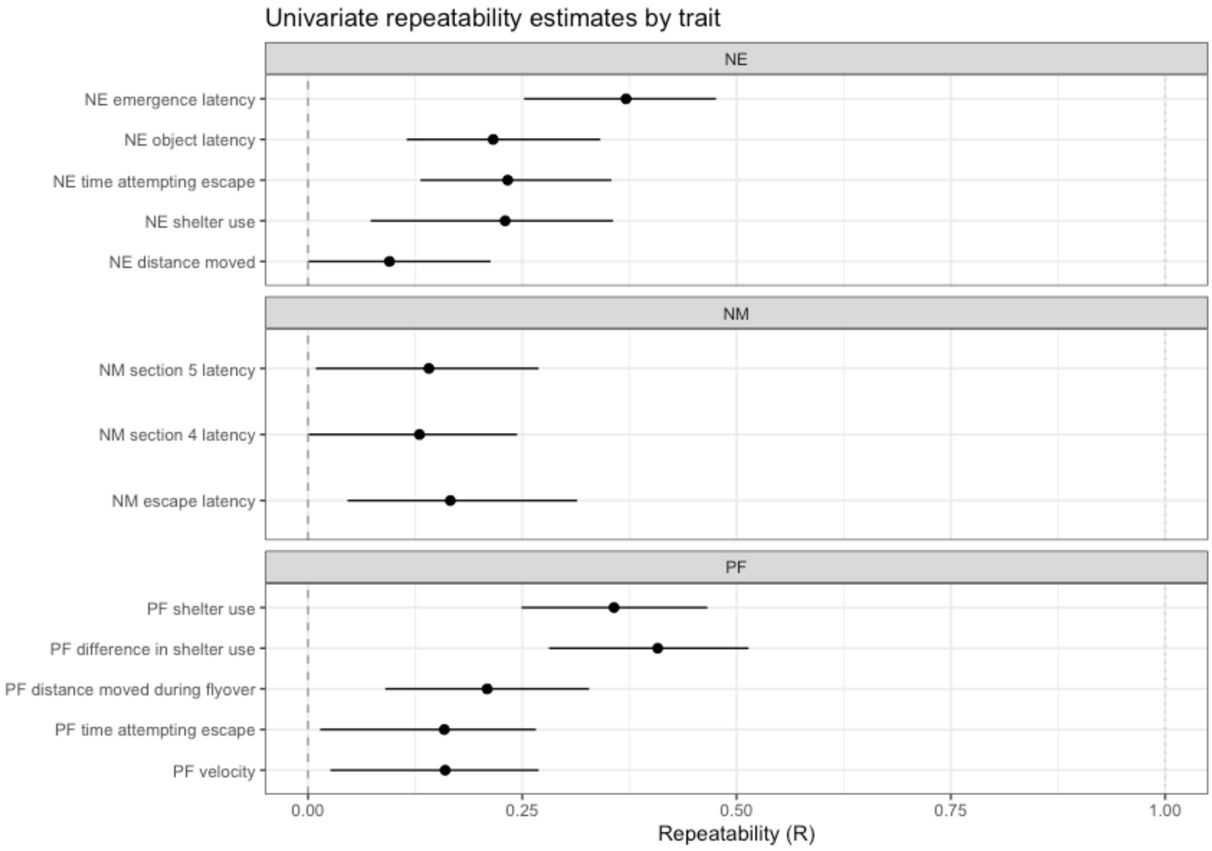

**Figure S4.** Univariate repeatability estimates (R) of the 13 behavioural measures included in the multivariate analysis. Novel Environment and Object assay is represented as NE, Novel Maze as NM, and Predator Flyover as PF.

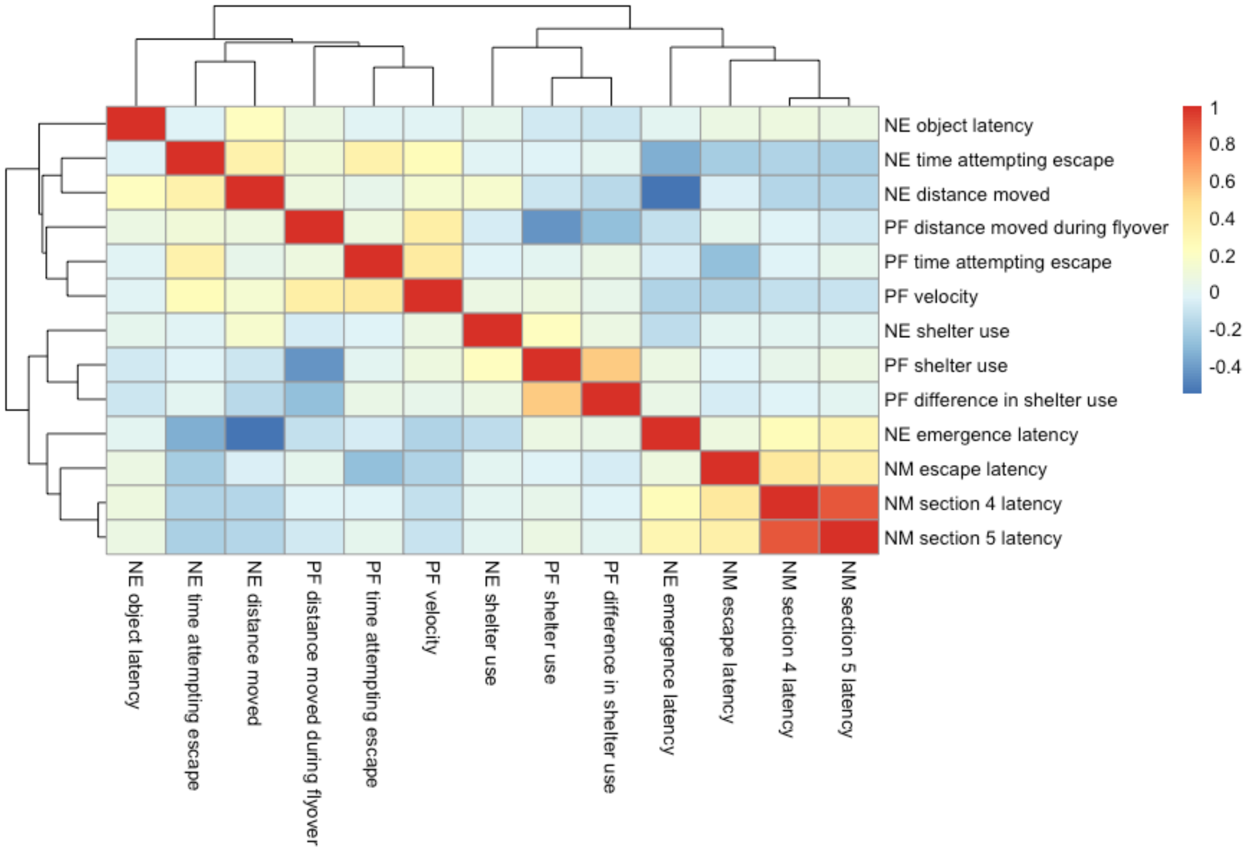

**Figure S5.** Covariance heatmap of behavioural measures included in the multivariate analysis to demonstrate the potential groupings of traits in the proposed models. Novel Environment and Object assay is represented as NE, Novel Maze as NM, and Predator Flyover as PF.

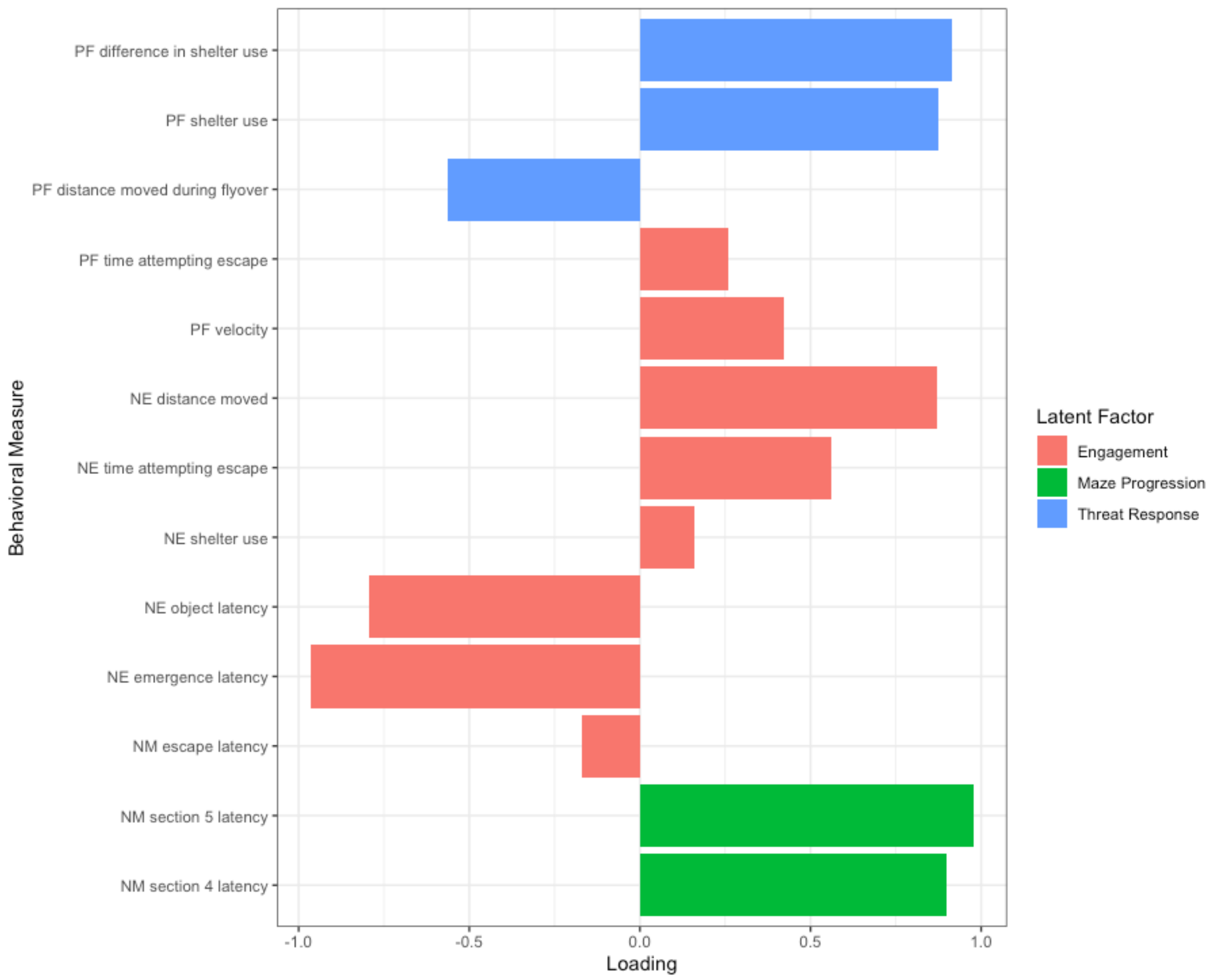

**Figure S6.** Magnitude (0-1) and direction (-/+) of behavioural measure loading onto the three latent factors identified from the best supported *a priori* model. Novel Environment and Object assay is represented as NE, Novel Maze as NM, and Predator Flyover as PF.
